# Context-dependent effects of antimicrobial coatings on microbial load and bacterial community composition on public high-touch surfaces

**DOI:** 10.64898/2026.02.05.703978

**Authors:** Harleen Kaur, Rohish Kaura, Kertu Tiirik, Marika Truu, Jaak Truu, Mati Kook, Dmytro Danilian, Vambola Kisand, Laman Mehraliyeva, Merja Ahonen, Meija Kivisaari, Jussi Tamminen, Aleksandr Semjonov, Angela Ivask

## Abstract

**Background:** Antimicrobial surfaces incorporated into high-touch public areas are used as a passive intervention to reduce surface microbial load and reduce the spread of infections. However, usually such surfaces lack proof of their antimicrobial activity in real use conditions and against a wider variety of microbes. This study evaluated the real-world performance of copper, TiO_2_-, silver- and quaternary ammonium compound (SiQAC)-based surfaces, which are commercially available and have proven antibacterial activity in lab tests. The surfaces were introduced to five study sites in diverse environments to collect data on bacterial load, community structure and taxonomic profile over several months.

**Results:** Copper surfaces introduced to shopping basket handles consistently exhibited the strongest antimicrobial performance, with significant reductions in aerobic bacterial counts and bacterial DNA, accompanied by clear shifts in microbial community composition. These shifts included reduction of several human-associated and opportunistic taxa and relative enrichment of environmentally resilient, stress-tolerant genera. TiO_2_-based photocatalytic coating reduced bacterial load in kindergarten tables under field conditions but did not significantly alter overall community structure. Silver-based surfaces on university campus tables showed minimal effects on microbial load and composition despite confirmed antibacterial activity in laboratory testing. Analogously to silver, SiQAC-based coating despite being active in lab conditions showed no decrease in bacterial load in real use conditions. When applied onto cafeteria and animal clinic tables SiQAC coating displayed context-dependent effects, with modest, genus-specific changes and increased richness in a high-contact cafeteria environment, but no significant impact in a low-biomass animal clinic setting. Viability based analysis revealed that on most of the surfaces a notable fraction of detected microbial DNA originated from non-viable cells.

**Conclusions:** This multisite field study demonstrates that the real-world performance of antimicrobially coated surfaces is strongly context dependent and cannot be reliably predicted from laboratory testing alone. Moreover, to understand the effect of antimicrobial coatings on surface microbial communities, real-use monitoring is needed.

## Introduction

High-touch communal surfaces act as dynamic microbial reservoirs that facilitate the exchange and transmission of microorganisms between natural sources, humans, and animals [1]. These surfaces may also serve as sinks for potential pathogens and act as sources for infectious diseases. Indeed, surface-mediated transfer is considered one of the main modes of interpersonal exchange of microbes and their related infectious diseases [2]. Antimicrobial surface coatings (AMC) have been suggested as passive means to decrease the bacterial residence time on surfaces and provide long-term protection against microbial surface colonisation [3]. AMC have gained attraction because they provide residual antimicrobial activity that can supplement conventional cleaning protocols and reduce the risk of the transfer of pathogens via high-touch surfaces to improve indoor hygiene [3,4]. In medical and veterinary settings, the use of AMC improves overall infection prevention and control of antimicrobial resistance or management of complex or mixed infections [5].

To verify the antimicrobial effect of an AMC, standardised laboratory testing should be carried out. Most traditionally, efficacy testing is performed following ISO 22196:2011, which foresees the incubation of surfaces with bacteria in a liquid environment and high humidity conditions [6]. However, such conditions differ significantly from most real-world scenarios, where surfaces are used under dry or semidry ambient conditions. To account for the latter, in 2023 a new ISO method (ISO 7581:2023) was introduced, which however has not been widely used yet in real surface (*in situ*) efficacy evaluations [7]. However, in general a trend indicating a drastic decrease in antimicrobial efficacy when moving from liquid high humidity test conditions to semi-dry or dry exposure conditions, exists [8,9,10]. Although ISO 7581 is a further development of ISO 22196 as it resembles more real-life use settings, both tests are performed in a laboratory.

However, more reliable and comprehensive data on the efficacy of antimicrobial surfaces could be obtained through field studies. To date, no standard methods for real-use monitoring of surface antimicrobial efficacy exist and a variety of custom methods have been used [11]. However, all those methods involve swabbing of a defined area of the surface, extracting the collected microbial cells using a buffer and either seeding the cells for aerobic colony counting or determining the cell number via other methods such as ATP extraction, qPCR, etc.) [12,13]. Such field monitoring studies have shown that selected antimicrobial surfaces indeed decrease the number of surface-residing bacteria [14, 15]. As expected, copper has proven to be the most active material, and its use in hospital environments has been shown to result in ∼ 1 log decrease in viable aerobic counts [16]. Similar ∼ 1 log decrease on copper surfaces have also been observed in kindergarten, retirement home and office building [17]. Interestingly, some studies have shown that the decrease in microbial counts due to the introduction of copper surfaces is use-case dependent [18–20]. For example, while ∼ 1 log reduction in the aerobic bacterial counts were observed on copper-coated preparation tables, shower trolleys, toilet seats and bed rails, whereas almost non-existent reduction has been observed on similarly coated door or cabinet handles [21,22]. In addition to copper, a limited number of real-life monitoring studies on silver and quaternary ammonium compound-based surfaces are available. Silver-containing surfaces have shown a real-life use effect by reducing surface bacterial counts by ∼ 1 log [23], similar to copper. Quaternary ammonium-based surfaces have demonstrated contradictory information, with one study showing high-level antimicrobial activity of more than 3 log reduction, whereas other studies showing no effect [24,25].

Microbial counts obtained from field samplings provide information on the efficacy of AMC against recoverable microbes but do not indicate which microbial taxa are affected. Yet the microbial community composition of the indoor surfaces plays a crucial role in creating a healthy living environment. Changes in indoor microbiome composition are likely to affect human well-being, especially if the composition of the indoor microbiome evolves towards pathogenicity and resistance [26,27]. It has been suggested that selection pressure on the indoor microbiome by antimicrobial selection may shift community structure, reduce taxonomic richness, and selectively enrich taxa with intrinsic or acquired tolerance to either the antimicrobials or antibiotics used [28]. One large-scale study carried out globally in several cities suggested that the spread of antibacterial resistance genes is increasing in human-built environments, which is, among other reasons, also likely due to the increasing use of antimicrobial compounds [29]. Nevertheless, studies examining changes in microbial community structure after the use of antimicrobial compounds are scarce. One earlier study looked at the community structure of marine biofilms on copper surfaces and demonstrated that copper exposure increased the proportion of Gram-negative facultative anaerobic microbes, that can survive across fluctuating redox conditions ranging from oxygen-rich surface layers to oxygen-limited biofilms [30]. A recent study conducted on copper surfaces in German schools and on the International Space Station did not find any differences in microbiome composition between copper and reference surfaces, which could be partially explained by the low biomass and quality of sequencing [31].

To more clearly understand how antimicrobial surfaces influence surface-associated microbial communities under routine use conditions, the AMC surfaces were deployed on basket handles in a hardware shop and on tables in a university campus, a kindergarten, cafeterias, and an animal clinic, followed by repeated field sampling. The objective of this study was to evaluate the real-world effects of four commercially available AMC (copper-based (Cu-T), a TiO₂-based photocatalytic coating (TiO₂-C), a silver-based (Ag-S), and a quaternary ammonium compound-based (SQ-C) surfaces) with previously demonstrated antibacterial efficacy. Specifically, we aimed to (i) quantify the effects of these AMC surfaces on surface-associated bacterial load using both culture-based and molecular approaches, (ii) assess how these AMC surfaces influence the composition, diversity, and structure of surface-associated microbial communities, and (iii) distinguish between viable and non-viable bacterial fractions to obtain a viability-resolved view of microbial persistence on treated surfaces. To achieve these aims, bacterial abundance was assessed using culture-based colony-forming unit (CFU) measurements and quantitative PCR, while community structure was characterised by sequencing the bacterial 16S rRNA gene. By integrating multisite field surface sampling with quantitative, taxonomic, and viability-based analysis, this study aims to provide an ecologically informed assessment of how AMC surfaces shape microbiota in real-life environments and to determine whether laboratory-demonstrated antimicrobial efficacy translates into meaningful effects under practical use conditions.

## Materials and methods

### Antimicrobial surface coatings

Table 1 presents the four different surface coatings that were tested for their antibacterial activity in laboratory conditions and for their effects on surface bacterial community at five sites across Finland and Estonia.

**Table 1.**
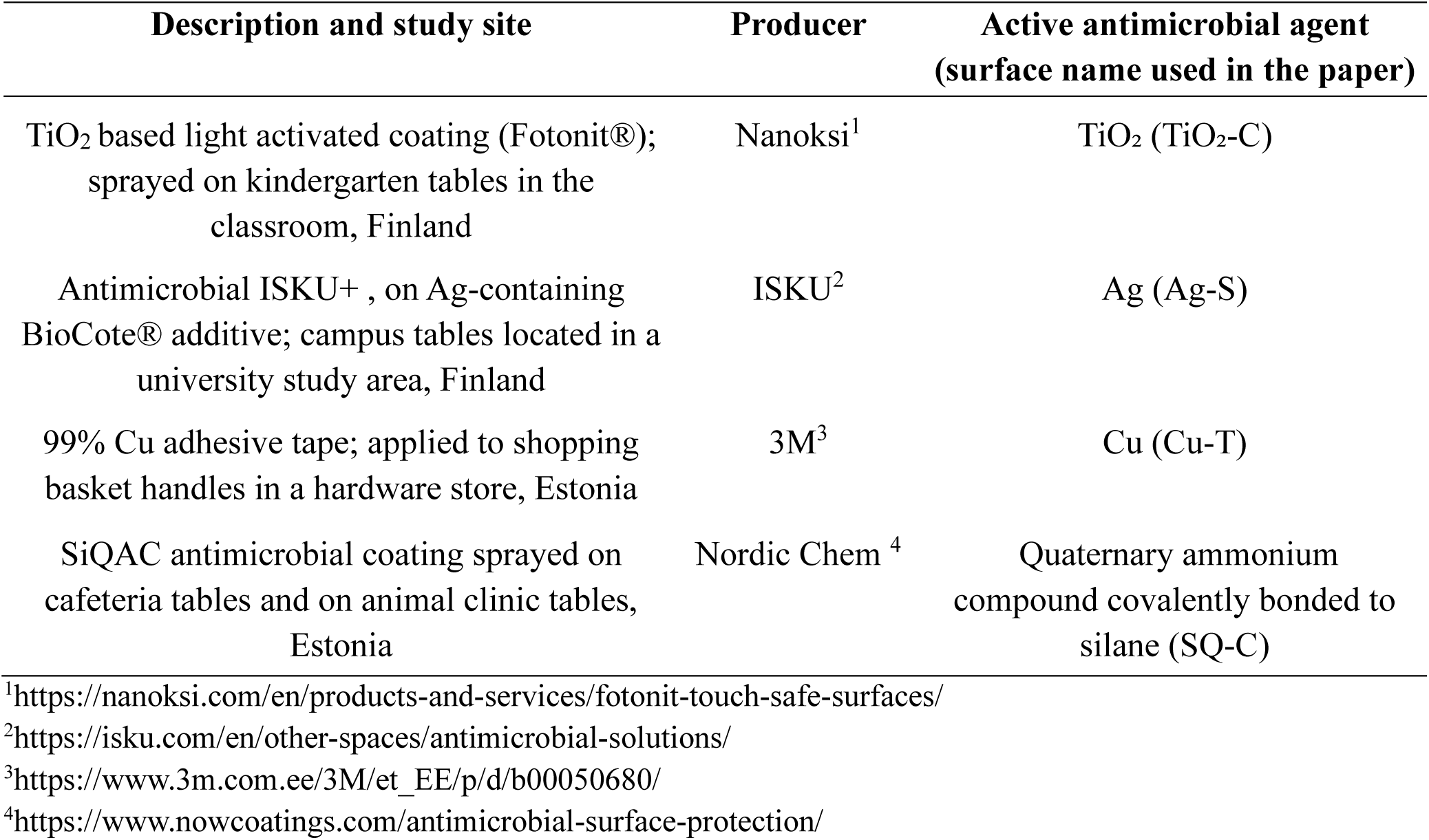
Description of antimicrobial surfaces used in the study.

### Laboratory analysis of antimicrobial efficacy of surface coatings

Prior to antibacterial testing, all four antimicrobial surface coatings listed in Table 1 were subjected to lab-scale physicochemical surface characterisation to verify the presence and integrity of the coatings after real-life–mimicking application. For laboratory analysis, TiO_2_-based (TiO_2_-C) and SiQAC-based (quaternary ammonium compound) (SQ-C) coatings were applied on 2.5×2.5 cm stainless steel (AISI 304) coupons by spraying and copper tape was adhered onto steel surface. Ag-based (Ag-S) surfaces were cut to 2.5×2.5 cm coupons. The coupons were examined using SEM for surface morphology, XPS for elemental composition, FTIR for chemical group identification, and for water contact angles for wettability estimation as detailed in materials and methods section in the supplementary. The ISO 22196:2011 method [9] was used to analyse the effects of Cu-T, Ag-S, SQ-C and stainless-steel surfaces against *Escherichia coli (E.coli)* ATCC 8739. The bacterial inoculum was prepared from fresh overnight cultures on LB agar plates (5 g/L yeast extract, 10 g/L tryptone, 5 g/L NaCl, and 15 g/L agar). The bacterial mass from the overnight plate was used to prepare cell suspension in 500-fold diluted nutrient broth (0.006g/L meat extract, 0.02g/L peptone, and 0.01g/L NaCl) at OD600 value of 0.005. A drop of 15 µL of bacterial suspension (ca 4.7×10^4^ CFU/surface or 1.5×10^4^ CFU/cm^2^) was pipetted onto 2.5×2.5 cm surfaces, covered with sterile 2×2 cm plastic foil and incubated at 35°C in a closed, humid environment for 1 h and 24 h. Following incubation, each surface was submerged into 10 mL of SCDLP (Soybean casein digest lecithin polysorbate-17 g/L casein peptone, 3g/L soybean peptone, 5 g/L NaCl, 2.5 g/L Na_2_HPO_4_, 2.5 g/L glucose, 1.0 g/L lecithin, and 7.0 g/L Tween 80) neutralising broth in a sterile 50 mL centrifuge tube. Surfaces were then vortexed for 30 seconds to detach the bacteria from the surface. The resulting wash-off suspension was serially diluted using phosphate-buffered saline (PBS: 8g/L NaCl, 0.2g/L KCl,1.44g/L Na_2_HPO_4_, 0.2g/L KH_2_PO_4_; pH 7.1) and plated on LB agar plates. The plates were incubated at 37 °C for 24 h and CFU were enumerated. Results are expressed as log_10_-transformed CFU counts per surface. The ISO 27447:2019 method was used to analyse the antibacterial efficacy of the light activated TiO_2_-C surfaces and stainless steel (negative control) against *E. coli* ATCC 8739 [32]. The bacterial inoculum was prepared from fresh overnight cultures on LB agar plates; the cell density was adjusted photometrically to achieve a target OD600 value of 0.04. The pieces of 2.5×2.5 cm of TiO_2_-C covered stainless steel were inoculated with 20 µL droplet of the bacterial inoculum (corresponding to approximately 2.5×10^5^ CFU/cm^2^) and covered with sterile 2×2 cm plastic foil. Surfaces were incubated under UVA (intensity at 315–400 nm spectral range 2.0 W/m^2^) or in the dark in Climacell EVO climate chamber (Memmert, Germany) at 90% relative humidity and 22°C for 4 h. After exposure, the surfaces were placed into 10 mL of SCDLP medium, cells were recovered by vortexing, plated for CFU counts as described above. All experiments were performed in three biological replicates.

### Preparation of surfaces for real-life study

The surfaces with antimicrobial coatings listed in Table 1 were prepared in accordance with the manufacturers’ guidelines. At least 30 samples were collected from each study site. SQ-C was applied by spray coating in two study sites. In the cafeteria, selected tables were fully coated with SQ-C, while uncoated tables within the same area served as non-coated controls. In the veterinary clinic, individual tables were divided into coated and uncoated halves, allowing each table to serve as its own internal non-coated control. Samples from SQ-C and control surfaces in both sites were collected over a two-month period following coating application. TiO_2_-C was sprayed onto all tables within one classroom unit of a kindergarten, whereas an adjacent, structurally identical classroom unit was left untreated and used as the non-coated control. Sampling was conducted over a 12-month period after coating installation to assess long-term performance under routine use conditions. Ag-S surfaces consisted of premanufactured antimicrobial tables obtained directly from the producer and installed in a university campus study area. Tables of identical design but without the antimicrobial Ag-S component were used as controls. Sampling was performed five years after installation, within the timeframe for which the manufacturer claims sustained antimicrobial activity of up to ten years. Copper surfaces (Cu-T) were installed as 99% copper adhesive tape applied directly to the handles of shopping baskets in a hardware store. Uncovered plastic handles served as controls. These Cu-T basket handles had been in continuous use for approximately five years prior to sampling, having been installed during the COVID-19 pandemic period. Surface samples were collected by swabbing a defined area of coated and corresponding control surfaces. Each surface sample consisted of the resulting swab eluate, which was further processed for all CFU counts, DNA extractions, and sequencing analysis as outlined in the workflow shown in Fig. 1.

**Figure 1.**
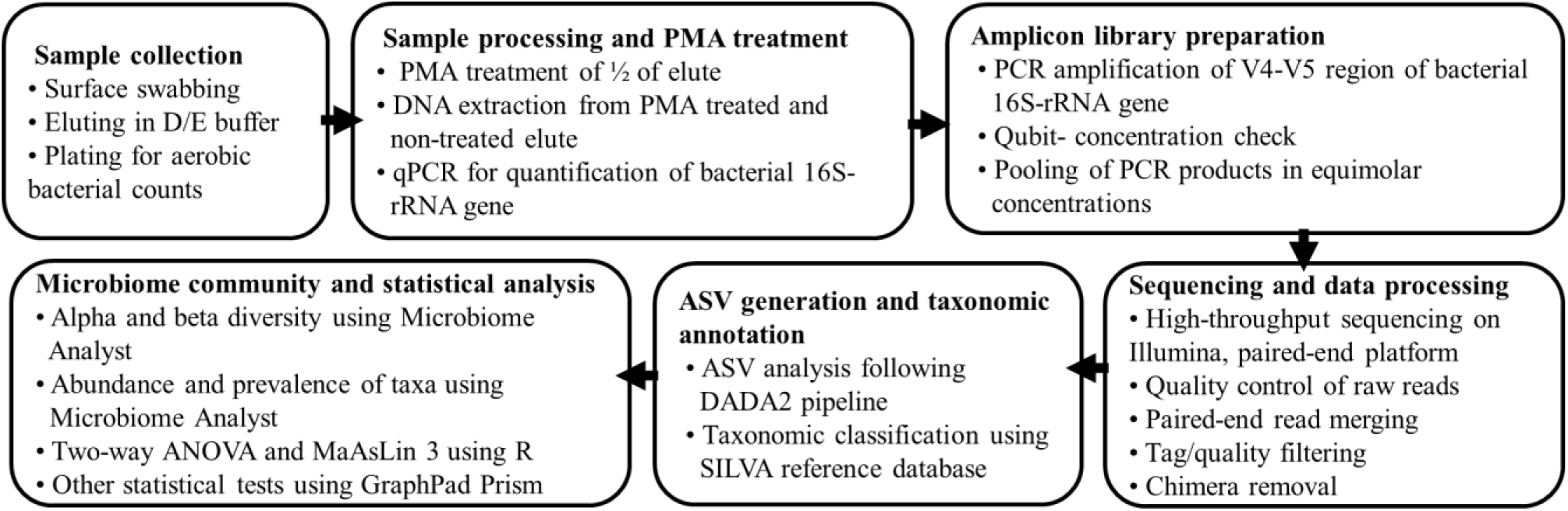
Flowchart of the workflow of the real-life microbial community analysis performed at the study sites. PMA: propidium monoazide used to distinguish DNA from dead or alive bacteria. ASV: amplicon sequence variant.

### Aerobic bacterial count on surfaces tested in real-life study

The surface samples were collected from a defined area in the study sites (Table 1) using sterile cotton swabs premoistened with 3.5 mL of Dey-Engley (D/E) neutralising diluent (BioTrading Benelux B.V., the Netherlands; composition: 10 g/L dextrose, 7 g/L lecithin, 6 g/L sodium thiosulfate, 5 g/L casein peptone, 5 g/L polysorbate 80, 2.5 g/L sodium bisulfate, 2.5 g/L yeast extract, 1 g/L sodium thioglycollate, and 0.02 g/L bromocresol purple) [33]. For the shopping basket handles, the entire copper-covered area was swabbed and as control area, plastic handles with the same size were swabbed. For tables, one premoistened swab was used to collect surface sample from a 200 cm² covered area. After surface swabbing, each swab was vortexed vigorously for 2 min to release microorganisms into the D/E diluent. Then, 1 mL of swab eluate was plated onto Compact Dry™ Total Count plates (Nissui Pharmaceutical Co., Ltd., Tokyo, Japan; nonselective medium with the redox indicator 2,3,5-Tripenyl Tetrazolium Chloride). The plates were incubated at 22°C for 5 days, and CFU were counted for estimating the aerobic bacterial load.

### DNA extraction and quantitative PCR from surface samples from real-life study

The remaining surface swab eluate of each sample was divided equally into two fractions and centrifuged to concentrate it to a final volume of approximately 200 µL. One fraction of the eluate was treated with propidium monoazide (PMA; PMAxx™ Dye; Biotium, Inc., Hayward, CA, USA) at a final concentration of 50 µM. PMA-treated surface samples were incubated in the dark for 10 min and subsequently exposed to a custom-built blue LED light source (465 nm) for 15 min to induce photoactivation and covalent crosslinking of PMA to DNA, according to the manufacturer’s instructions. PMA penetrates cells with compromised membranes and, upon photoactivation, binds to DNA, thereby preventing the amplification of DNA from dead or damaged cells. Therefore, only signal from alive bacteria could be amplified after PMA treatment. The other fraction of each the eluate was left untreated (non-PMA), and it served as a measure of total bacterial taxa [34]. Genomic DNA was extracted from all PMA-treated and non-PMA surface samples using the DNeasy PowerSoil Pro Kit (Qiagen, Hilden, Germany), following the protocol provided by the manufacturer. In addition, eluate of an empty cotton swab (not used for sampling) was extracted with the DNA kit. The extracted DNA was used to quantify the number of bacteria on surfaces by bacterial-specific 16S rRNA gene copy numbers using quantitative PCR (qPCR). The qPCR was carried out using Rotor-Gene Q real-time PCR cycler (Qiagen, Hilden, Germany) as described by Truu et al. 2020 [35] using SYBR Green qPCR Mix (Thermo Fisher Scientific Inc.), optimised concentrations of primers Bact517F (5′-GCCAGCAGCCGCGGTAA-3′) [36], and Bact1028R (5′-GACARCCATGCASCACCT-3′) [37], and following qPCR programme: 50°C 2 min, 95°C 10 min; 35 cycles: 95°C 30 s, 60°C 45 s, 72°C 45 s. All reactions were performed in triplicate, and each qPCR run included three no-template negative controls. The quantification data were analysed with the LinRegPCR program v.2020.0 (Ruijter et al., 2009) and the 16S rRNA gene copy numbers were calculated by estimating the fold difference between each sample and multiple data points of the standard curve, as described by Nõlvak et al. (2016) [38]. The data are presented as gene copy numbers per cm^2^ of surface (copies/cm^2^).

### 16S rRNA gene amplicon preparation, sequencing and sequence analysis

Bacterial community taxonomic profiling was performed using Illumina® NovaSeq 6000 sequencing of combinatorial sequence-tagged PCR products, according to the protocol described by Truu et al. 2020 [35]. The universal primers Bac515F (5′-GTGYCAGCMGCCGCGGTAA-3) and Bac926R (5′-AGTCAGCCAGCCGYCAATTYMTTTRAGTTT-3’) were used to target the V4-V5 hypervariable region of the 16S rRNA gene [35, 39]. DNA was amplified using the following programme: 98°C 30 s; 25-35 cycles: 98°C 10 s, 60°C 30 s, 72°C 15 s; final extension 72°C 8 min using thermal cycler Mastercycler X50 (Eppendorf AG, Hamburg, Germany). The concentration of each PCR product was determined using Qubit™ 4 Fluorometer (Thermo Fisher Scientific Inc., Waltham, MA, USA). For sequencing library preparation, PCR products from all individual surface samples were pooled in equimolar amounts. The mixture was purified and concentrated using the NucleoSpin® Gel and PCR clean-up Mini Kit (Macherey-Nagel GmbH & Co. KG, Düren, Germany), following the manufacturer’s protocol. Sequencing was performed by Novogene (Planegg, Germany) and the process involved pooling and end-reparation, A-tailing of PCR products and further ligation with Illumina adapters. Libraries were sequenced on Illumina® NovaSeq 6000 platform. In parallel with eluates from surface swab eluates. To assess the potential background contamination associated with sampling, DNA extraction and PCR amplification, sequencng library was also prepared from empty swab control (Table S1). As an additional control, sequencing library was alos prepared from a mock community (Zymo mock, product D6300) (Table S2).

The obtained raw paired-end amplicon sequencing reads were processed using QIIME2 [40]. Sequence quality control, trimming, denoising, paired end read merging, and chimera removal were performed using the DADA2 plugin [41]. Forward and reverse reads were truncated based on quality profiles to remove low-quality regions and sequencing adapters prior to denoising. DADA2 learned run-specific error models and inferred amplicon sequence variants (ASV) with single-nucleotide resolution. Chimeric sequences were identified and removed using the consensus method implemented in DADA2. The resulting high-quality ASV table and representative sequences were used for downstream analyses.

Taxonomic assignment of ASV was conducted in QIIME2 using a naïve Bayes classifier trained on the SILVA reference database version 138.2 [42]. Bacterial community data analysis was performed using Microbiome Analyst 2.0 software [43]. Initial preprocessing involved the removal of low-quality ASV. To ensure comparability across surface samples, data normalisation was performed using Total Sum Scaling (TSS), which adjusts for differences in sequencing depth. ASV were further refined by excluding those with fewer than four reads and those present in less than 10% of the total, thereby minimising the influence of rare taxa. Additionally, ASV exhibiting low variance, specifically those within the bottom 5% of the interquartile range (IQR) of variance were removed. Raw data used for the analysis can be found in the depository file with the accession number PRJEB106797 in ENA database. To analyse the diversity, richness and evenness of the communities, alpha diversity was calculated based on Observed ASV and Shannon index. For beta diversity analysis, ASV abundances were CLR-transformed, and community structure was assessed using principal component analysis (PCA). Taxonomic relative abundance of plots at genus level were generated in MicrobiomeAnalyst using the “merged” option, whereby sequencing reads were aggregated across all surface samples within each group and re-normalized to relative abundance, resulting in a single composite community profile per group.

Potential multivariable associations between genus-level microbial abundances and surface type (antimicrobially coated surface and respective control surface) were explored using MaAsLin3 (microbiome multivariable association with linear models, v3), while adjusting for PMA treatment (to separate the presence of total and alive bacterial taxa). Genus-level feature tables derived from 16S rRNA amplicon sequencing were filtered prior to analysis based on minimum abundance (≥1 × 10⁻⁴) and prevalence (≥10% of surface samples). Control surfaces and non-PMA surface samples were used as reference groups. We initially ran the models with interaction terms, but no genera showed significant associations. Therefore, we proceeded with a main-effects–only model to retain statistical power and interpretability.

### Statistical analysis

Two-way ANOVA followed by Dunnett’s multiple comparisons test and Fisher’s test at the p-value (0.05) were used for evaluating statistical significance of results from antibacterial test. Kruskal-Wallis test with Dunn’s post-hoc correction were used for comparisons of bacterial load on control surfaces across different study sites. Two-way ANOVA using Yeo-Johnson transformed data was used to detect significant differences between groups in microbial plate counts, bacterial 16S rRNA gene copy numbers, and alpha diversity indices (observed ASV and Shannon index) [44]. Statistically significant differences in microbial composition on surface samples from different study sites were revealed by differential abundance analyses using Linear Discriminant Analysis Effect Size (LEfSe) with a LDA score ≥ 2 and with significance threshold set at p < 0.05. Beta diversity differences were evaluated using permutational multivariate analysis of variance (PERMANOVA). Statistical significance for MaAsLin 3 results was assessed using Wald tests, and p-values were adjusted for multiple testing using the Benjamini–Hochberg false discovery rate (FDR) procedure. Associations with an FDR-adjusted q-value < 0.10 were considered statistically significant. Effect sizes (β coefficients) and 95% confidence intervals were discussed at the genus level and visualised using forest plots. Statistical analyses were performed using GraphPad Prism version 10.6.0 (GraphPad Software, Boston, MA, USA) and R version 4.4.1 (R Foundation for Statistical Computing, Vienna, Austria).

## Results

### Physico-chemical characteristics of the studied surfaces

To identify the presence and properties of the AMC surfaces, lab-scale characterisation of those coatings on stainless steel was performed. The detailed results are shown in supplementary Fig. S1-S4. SEM revealed that both TiO_2_-C and SQ-C coatings formed uniform layers (Fig. S1 B, E). A closer view of TiO_2_-C revealed the presence of 10-20 nm sized nanoparticles, which were suggested to be TiO_2_ particulates (Fig. S1 F). The presence of TiO₂ in the TiO_2_-C was further confirmed by Ti 2p X-ray photoelectron spectroscopy (Fig. S2). To provide more detailed identification and characterisation of SQ-C, Fourier-transform infrared spectroscopy analysis was performed, which confirmed the presence of quaternary ammonium functional groups in that coating (Fig. S3 A). Cross-sectional SEM imaging estimated the thickness of the SQ-C layer to be approximately 170–210 nm (Fig. S3 B). Cu-T surface had metal-like appearance under SEM (Fig. S1 C) and Ag-S showed a rather smooth surface (Fig. S1 D). The elemental contents of the Cu-T and Ag-S surfaces were not analysed, and the description of producer (Table 1) was trusted. Contact angle measurements showed that Cu-T, TiO_2_-C, and SQ-C surfaces have contact angles slightly above 90°, indicating mildly hydrophobic surface properties, while Ag-S and its corresponding control surface were hydrophilic with contact angles below 90° (Fig. S4). Compared with control surfaces none of the antimicrobial surface coatings changed the surface contact angle significantly (p > 0.05) suggesting that wettability characteristics of control and coated surfaces were similar and their antibacterial properties could be compared.

### Laboratory scale antimicrobial efficacy testing of the studied surfaces

All the surfaces were screened for their antibacterial properties in lab tests using *E. coli* ATCC 8739. The Ag-S, Cu-T, and SQ-C surfaces were evaluated using ISO 22196 method [9], with stainless steel used as the noncoated control surface. After 1 h of exposure, Cu-T surfaces achieved a 3 log₁₀ reduction in viable *E. coli* counts, whereas the SQ-C surfaces showed a 1.2 log₁₀ reduction (Fig. 2A). The Ag-S surfaces exhibited no measurable antibacterial activity at this early timepoint. The moderate CFU reduction observed in case of SQ-C and no CFU reduction measured in case of Ag-S at 1 h suggest that for those surfaces extended exposure is needed for maximal efficacy. After 24 h of exposure in lab tests, all the tested surfaces demonstrated a maximum 3 log_10_ reduction in viable counts (Fig. 2A) and therefore based on lab test results, could be expected to show efficacy also in real-life study which was carried out over several months.

**Figure 2.**
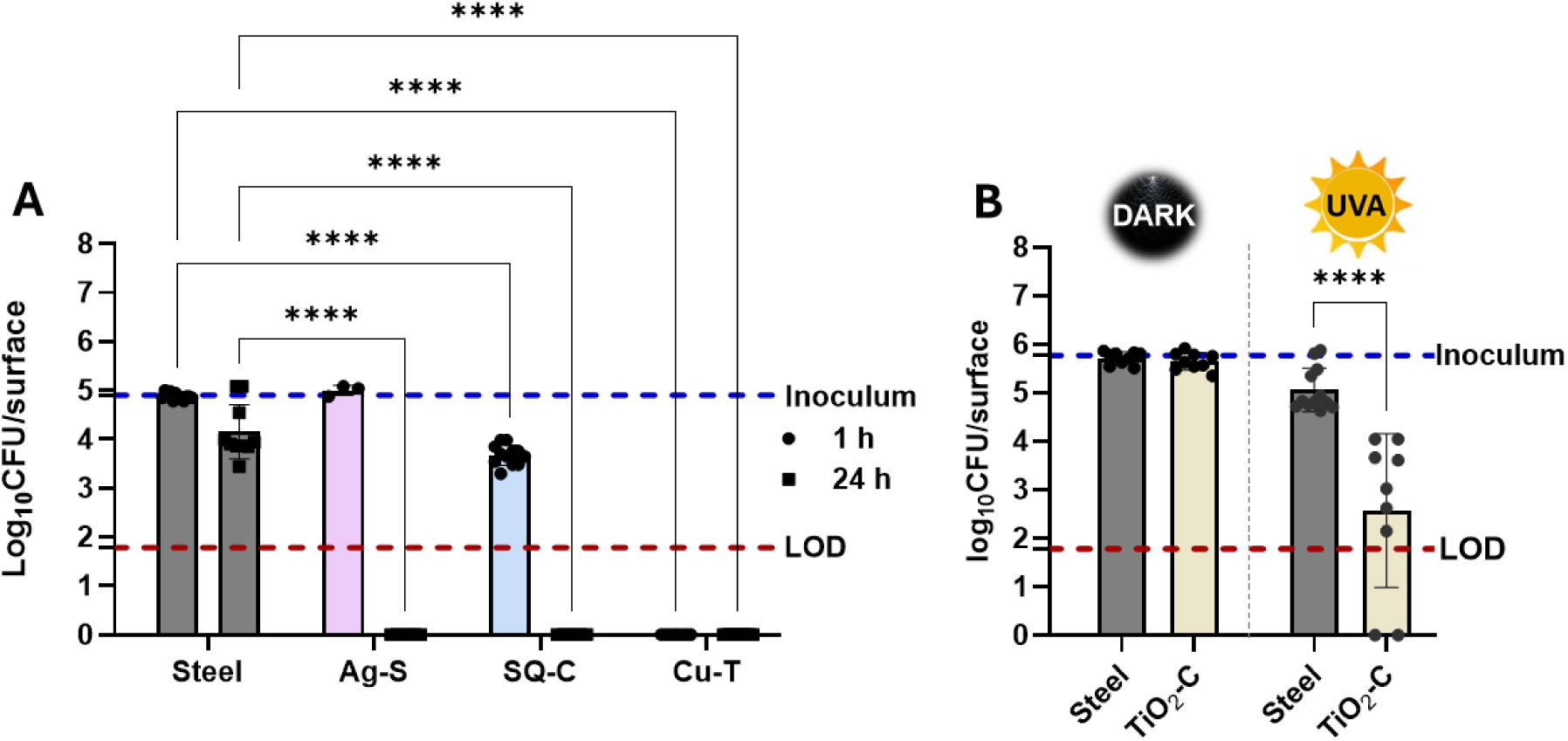
Antibacterial activity of Ag-based (Ag-S), SiQAC-based (SQ-C), and copper-based (Cu-T) surfaces in ISO 22196 test (A) and UVA driven antibacterial effect of TiO_2_-based (TiO_2_-C) surfaces in ISO 27447 test (dark incubation shown for reference) (B) against *E. coli* ATCC 8739. Each point represents the mean log₁₀ CFU per surface, error bars indicate standard deviation. The red dotted line denotes the limit of detection (LOD), and blue dotted line corresponds to the initial bacterial load (inoculum). Two-way ANOVA followed by Dunnett’s multiple comparison test(A) and uncorrected Fisher’s LSD(B) were used for the comparisons of the tested surfaces from their respective controls. Statistically significant differences are shown as **** (p<0.0001).

Due to the photocatalytic mode of action of TiO₂-based (TiO_2_-C) surfaces, the antimicrobial efficacy assessment in lab followed ISO 27447 protocol [32]. After 4 h under UVA TiO_2_-C surfaces resulted in a 2.5 log₁₀ reduction in viable *E. coli* counts as compared to the control (Fig. 2B). Therefore, in the presence of sufficient UVA, this surface could also be expected to exhibit antibacterial efficacy within the duration of the real-life study.

### Bacterial load and community structure of control surfaces across different study sites

The study sites for real-life efficacy and surface microbiome study included different types of environmental conditions, microbial exposure risks, and patterns of surface-human interaction: tables in a kindergarten classroom and university campus in Finland, and cafeteria, animal clinic, and shopping basket handles in a hardware store in Estonia (Table 1). Across all the study sites, the average aerobic bacterial load on the control surfaces was below the standard benchmark of 5 CFU/cm² (Fig. 3A) which is generally regarded as indicative of good hygiene practices [45]. However, some differences were observed in bacterial load in the study sites, which may be attributed to different cleaning patterns and surface usage conditions. Shopping basket handles exhibited the highest bacterial loads, averaging 4.1 CFU/cm², followed by kindergarten tables (2.9 CFU/cm²) and cafeteria tables. The bacterial load of the campus and animal clinic tables was relatively low. Quantification of the 16S rRNA gene in the study sites was, in general, in agreement with CFU data. For instance, on average, the high plate counts and gene copy numbers were recorded on shopping basket handles (Fig. 3B). However, the animal clinic tables, which had lower bacterial plate count, than tables in kindergarten, campus and cafeteria, the 16S rRNA gene abundances exceeded the abundances on the other three sites tables (Fig. 3B). This discrepancy could be due to differing fraction of cultivable bacteria at different study sites.

**Figure 3.**
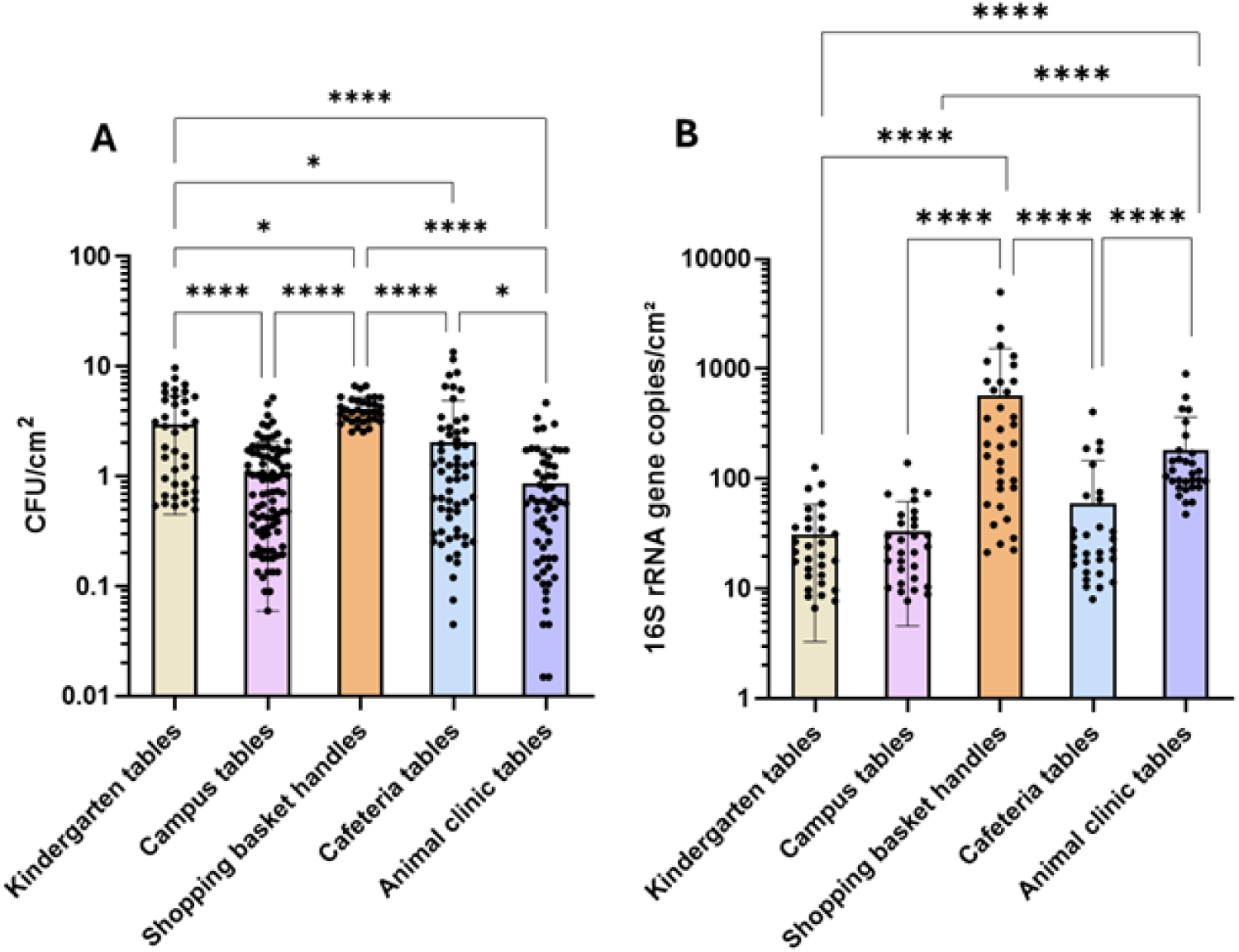
Bacterial counts (CFU/ cm²) (A) and 16S rRNA gene copy numbers (B) on the surfaces of five study sites. The bars represent mean values ± standard deviations; individual data points are shown as black dots. Statistically significant differences between groups were determined using Kruskal-Wallis test followed by Dunn’s multiple comparisons tests and shown as * (p<0.05) and **** (p<0.0001).

Community analysis revealed that different microbial taxa were present at the different study sites. Alpha diversity analysis based on either the number of observed ASVs, which reflects microbial richness, or Shannon index, which accounts for both richness and evenness, revealed significant differences across the sampled sites. (Fig. 4A, 4B). Compared with all other environments, basket handles showed the highest alpha diversity, while in the tables of animal clinic, this parameter was the lowest. Beta diversity analysis based on PCA further supported these findings by revealing distinct clustering of microbial communities according to environment (Fig. 4C). Besides, PERMANOVA showed a significant effect of study site on microbial community composition (F = 48.29, R² = 0.57, p = 0.001). These statistics represent overall differences between all the study sites and do not correspond to pairwise comparisons. PCA revealed that the bacterial community on basket handles formed a well-separated and distinct cluster, whereas differences among bacterial communities from cafeteria, animal clinic, kindergarten, and campus tables were less pronounced. Bacterial communities associated with animal clinic tables showed slight separation from campus tables but overlapped with communities on kindergarten and cafeteria tables (Fig. 4C).

**Figure 4.**
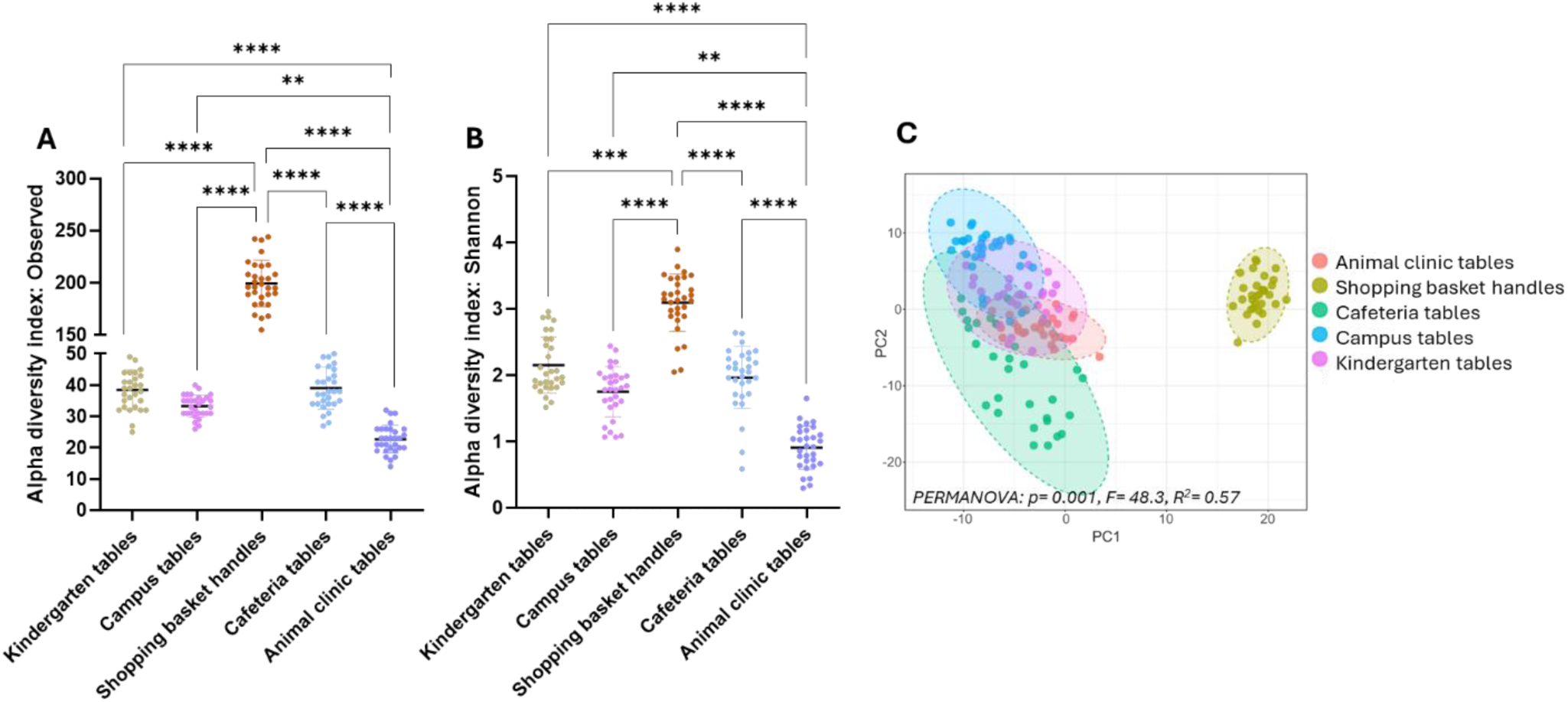
Results of alpha and beta diversity analysis of the bacterial community on control surfaces from the five study sites. For alpha diversity index based on Observed ASV (A) and Shannon (B) results of individual samplings with mean values ± standard deviation are shown. Comparisons between sampling sites are based on Kruskal-Wallis test followed by Dunn’s multiple comparisons tests, statistically significant differences are shown (**p < 0.01, ***p < 0.001, ****p < 0.0001). (C) PCA plot showing clustering of the study sites based on bacterial community structure, with statistical significance assessed by permutational multivariate analysis of variance (PERMANOVA).

The bacterial communities of shopping basket handles were dominated by *Staphylococcus* (36.4% of all detected genera) and *Streptococcus* (8.2%), while cafeteria surfaces showed high proportions of *Pseudomonas* (36.1%) and *Rhodococcus* (18.2%), and kindergarten tables were dominated by *Rhodococcus* (36.4%) and *Arthrobacter* (12.4%) (Fig.5). The animal clinic exhibited a strong dominance of *Pseudomonas* (77.2%), whereas campus tables were primarily characterised by *Rhodococcus* (38.0%), *Arthrobacter* (13.7%), and *Pseudomonas* (9.2%). Across all study sites, genera such as *Rhodococcus*, *Pseudomonas*, and *Arthrobacter, Staphylococcus, Sphingomonas, Streptococcus* and *Cutibacterium* emerged as core members of the microbiome (Fig. S5). Although some of these species are commonly associated with soil and natural environment, earlier studies have shown the presence of all of those in human-related environments [46]. Members of genus *Pseudomonas, Staphylococcus, Sphingomonas, Streptococcus* and *Cutibacterium* have been shown on a variety of high-touch surfaces and built environment in general [47–50]. *Arthrobacter* and *Rhodococcus* have been related with touch surfaces less frequently but have been found in long-term care facilities [51], in subway surfaces [46,48, 52], and as a part of air microbiome [51,48].

**Figure 5.**
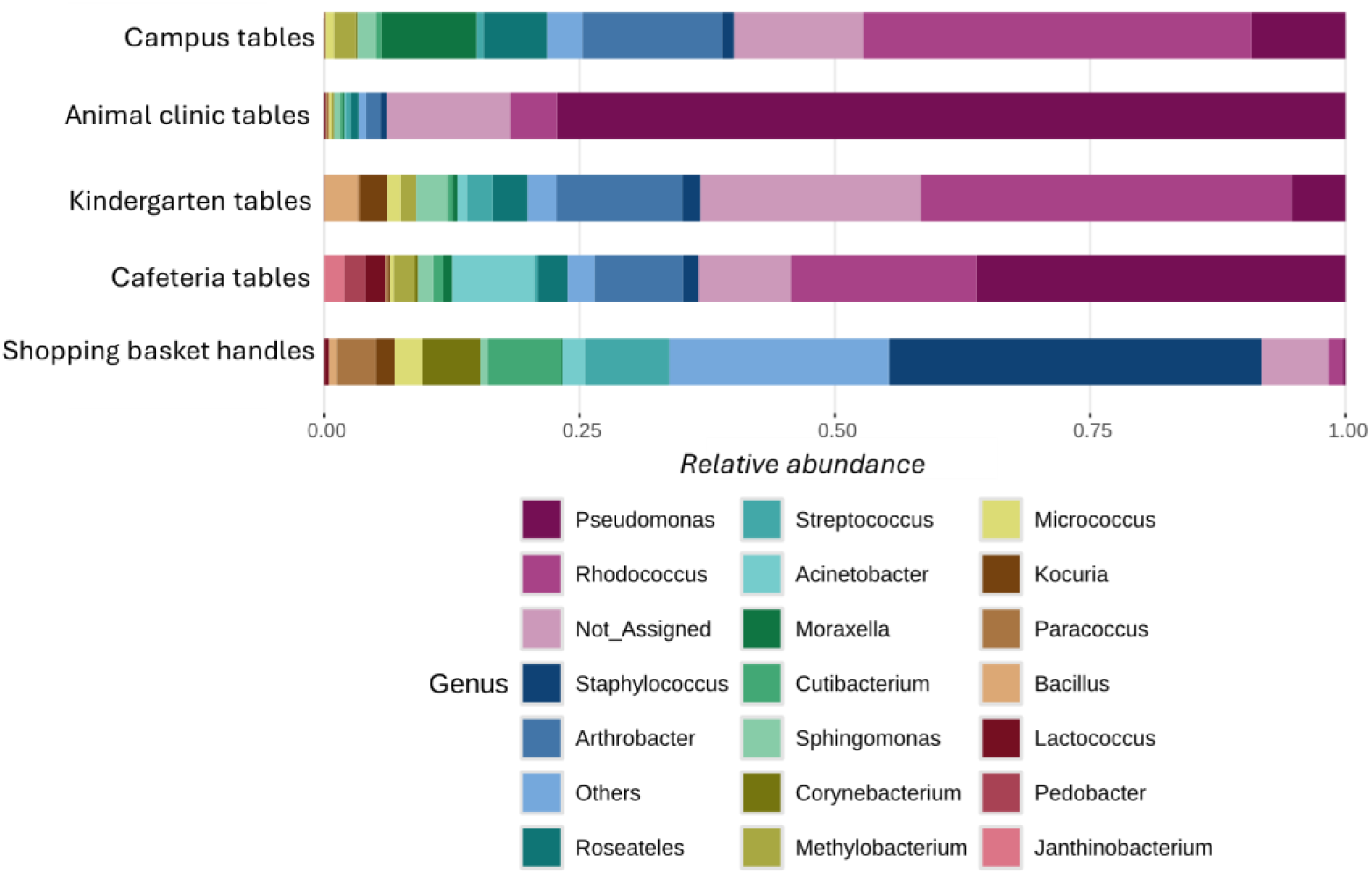
Relative abundance of top 20 bacterial genera across the five study sites. Stacked bar charts represent the merged proportional composition of bacterial genera on control surfaces of each site. The relative abundance of bacterial genera of individual surface sample as shown in Fig. S6.

LEfSe analysis also indicated differences between the study sites (Fig. 6) indicating enrichment of *Staphylococcus, Cutibacterium, Streptococcus, Corynebacterium, Paracoccus, Micrococcus and Kocuria* (LDA >4) in shopping basket handles, *Pedobacter*, *Janthinobacterium*, *Acinetobacter*, *Roseateles*, and *Methylobacterium* (LDA >4.0) in cafeteria tables, *Pseudomonas* (LDA >4.0) and *Lactococcus* in animal clinic, *Arthrobacter*, *Bacillus*, *Sphingomonas*, *Dermacoccus* and *Microbacterium* in kindergarten and *Rhodococcus, Moraxella*, *Roseateles*, and *Acinetobacter* in campus tables. These results showed that although some genera repeated in different environments, clear site-specific differences appeared that likely resulted from user specificity. The genera enriched in shopping basket handles most resembled human skin-related microbiome (genera *Staphylococcus, Cutibacterium, Streptococcus, Micrococcus*). On cafeteria tables the enriched genera were mostly related with natural environment such as soil and water and in general, the same with some exceptions (e.g. *Dermacoccus* in kindergarten), can be concluded for kindergarten and campus tables. Animal clinic showed a clear enrichment of *Pseudomonas*, which is a genus with many natural, clinical and industrial environments.

**Figure 6.**
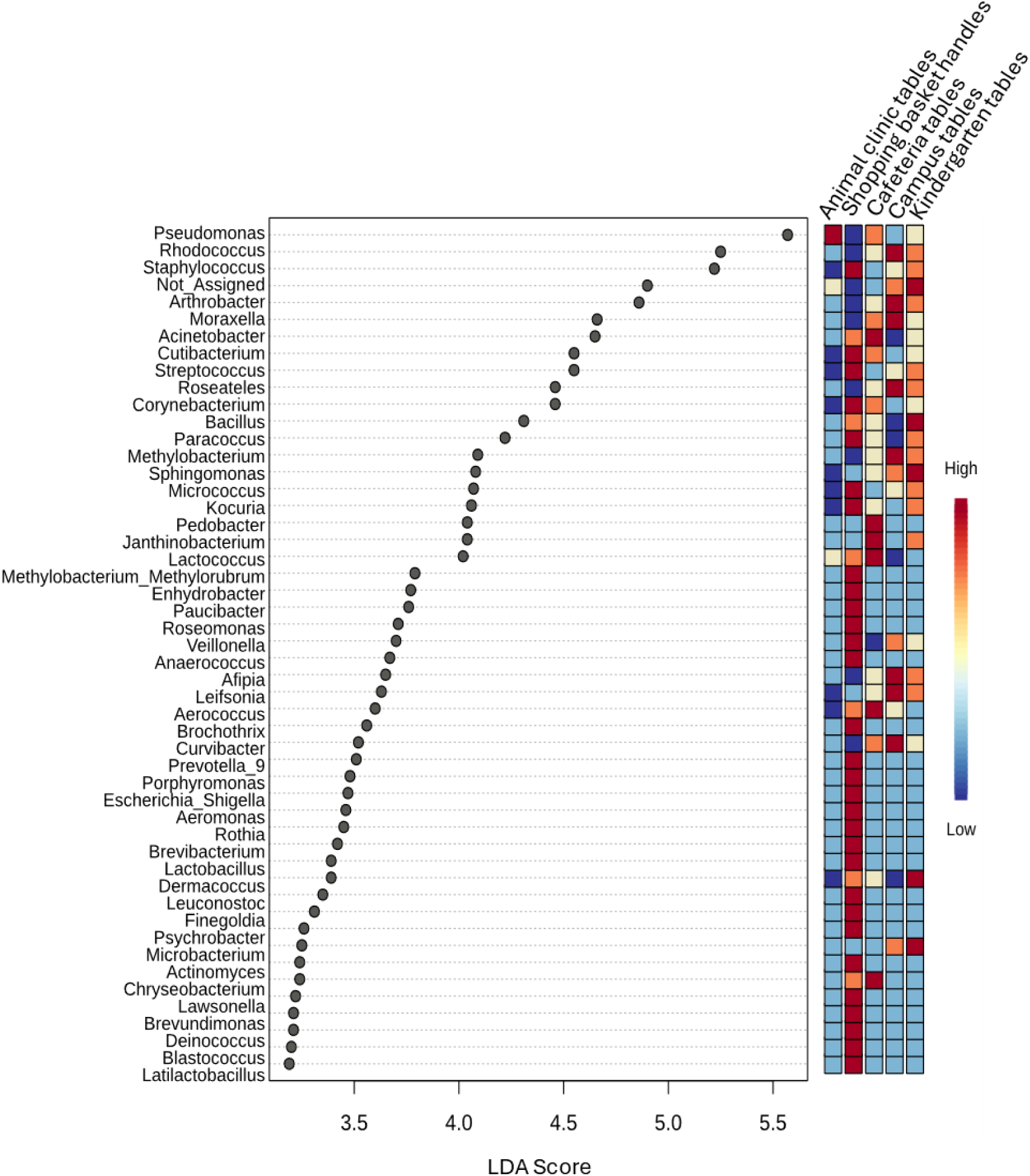
Plot showing the top 50 bacterial genera significantly associated with different sampling study sites according to the LEfSe analysis. Genera with LDA score ≥ 2 and p < 0.05 are shown on the y-axis, with effect size on the x-axis. The heatmap displays relative abundance across five study **sites**.

### Effect of copper-based coatings on microbiota of basket handles

#### Microbial load and diversity

Comparison of aerobic bacterial counts of surface samples from Cu-T and control (non-copper) basket handles showed that introduction of Cu-T, significantly reduced the bacterial load on basket handles (p < 0.0001; Fig. 7A). Consistent with these results, 16S rRNA gene copy numbers were also significantly reduced on Cu-T (p < 0.001; Fig. 7B). Comparison between 16S rRNA gene copies on AMC and control surfaces was carried out after PMA treatment, which is claimed to leave only DNA from living microbes available for PCR amplification [53] and has been used to discriminate between alive and dead cells in several studies. Our results showed that, even after PMA treatment, 16S rRNA copies on control and Cu-T basket handles differed (p < 0.001; Fig. 7E).

**Figure 7.**
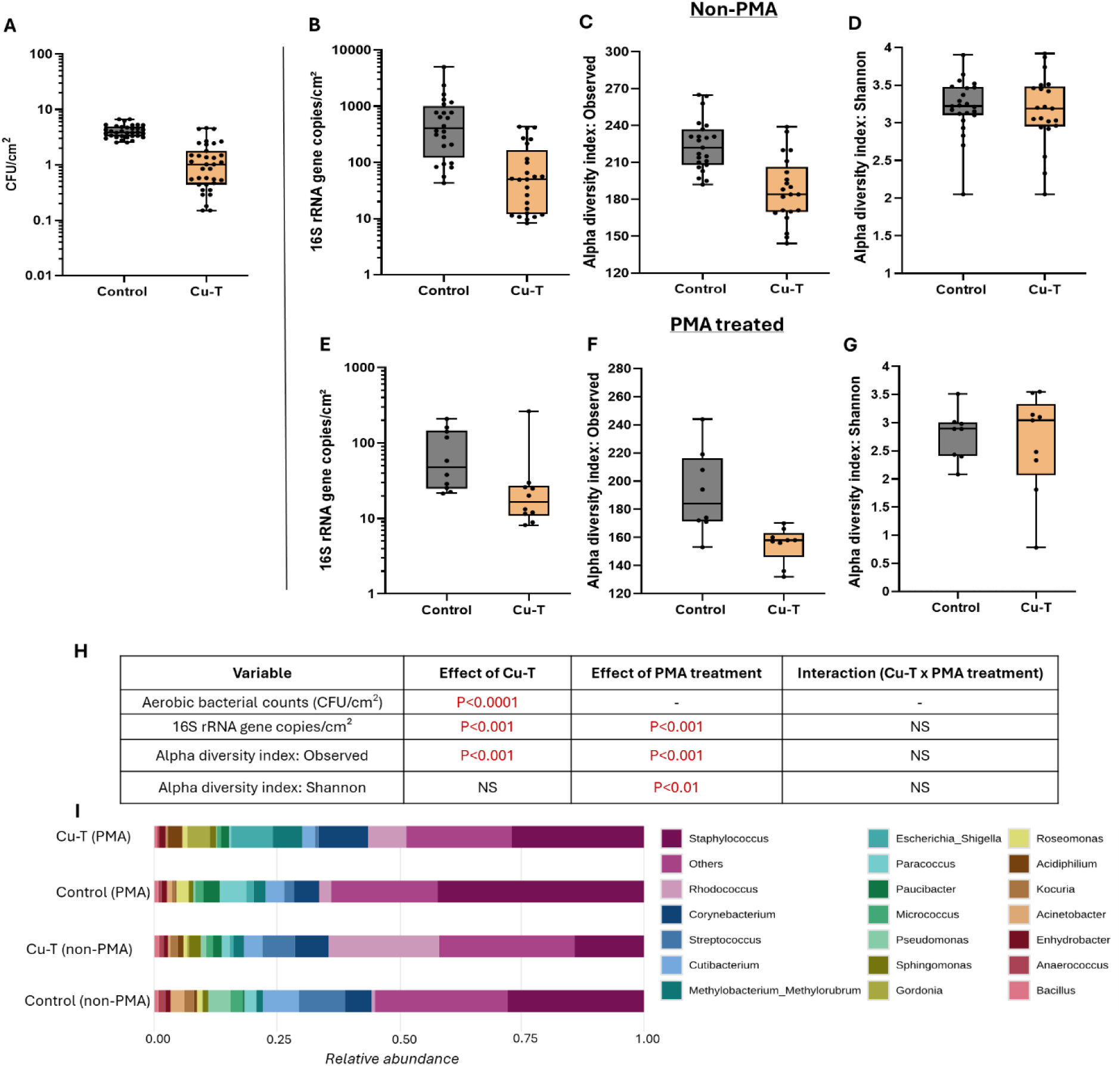
Comparison of bacterial load and community basket on copper (Cu-T) and control basket handles. Aerobic bacterial counts (CFU/cm²) (A), 16S rRNA gene copies per cm² for non-PMA(B) and PMA(E), Observed ASV for non-PMA (C) and PMA(F), Shannon diversity index for non-PMA(D) and PMA(G) treated surface samples. Box-and-whisker plots show the median (center line), interquartile range (25th–75th percentiles; box), and minimum–maximum values. Individual data points are overlaid as dots. (H) Summary of statistical tests evaluating the effects of Cu-T on CFU counts, 16S rRNA gene abundance, observed ASV, and Shannon diversity. One-way ANOVA was applied to CFU data, and two-way ANOVA was applied to Yeo–Johnson–transformed data to test the effects of Cu-T, PMA treatment, and their interaction. NS = not significant (p ≥ 0.05). (I) Relative abundance of the top 20 genera across Cu-T and control surfaces for PMA non-treated and treated datasets; stacked bars represent merged proportional composition of bacterial genera from all surface samples per group with colors indicating individual genera. Relative abundance profiles of genera for individual surface samples are presented in Fig. S7.

Alpha diversity analysis revealed a significant reduction in richness based on Observed ASV counts on Cu-T surfaces as compared with control surfaces irrespective of PMA treatment or no treatment (p < 0.001; Fig. 7C, 7F). In contrast, Shannon diversity did not differ between Cu-T and control surfaces for either non-PMA or PMA-treated surface samples. (Fig. 7D, 7G). This suggests that while Cu-T reduced the number of detectable taxa, the relative abundance distribution among the remaining taxa was largely preserved, meaning evenness was not substantially affected.

Beta diversity analysis revealed significant differences in microbial community composition between Cu-T and control surfaces. PCA showed clear statistically significant differences between Cu-T and control surfaces, which was evident both in case of non-PMA (Fig. 8A) and PMA-treated surface samples (Fig. 8B).

**Figure 8.**
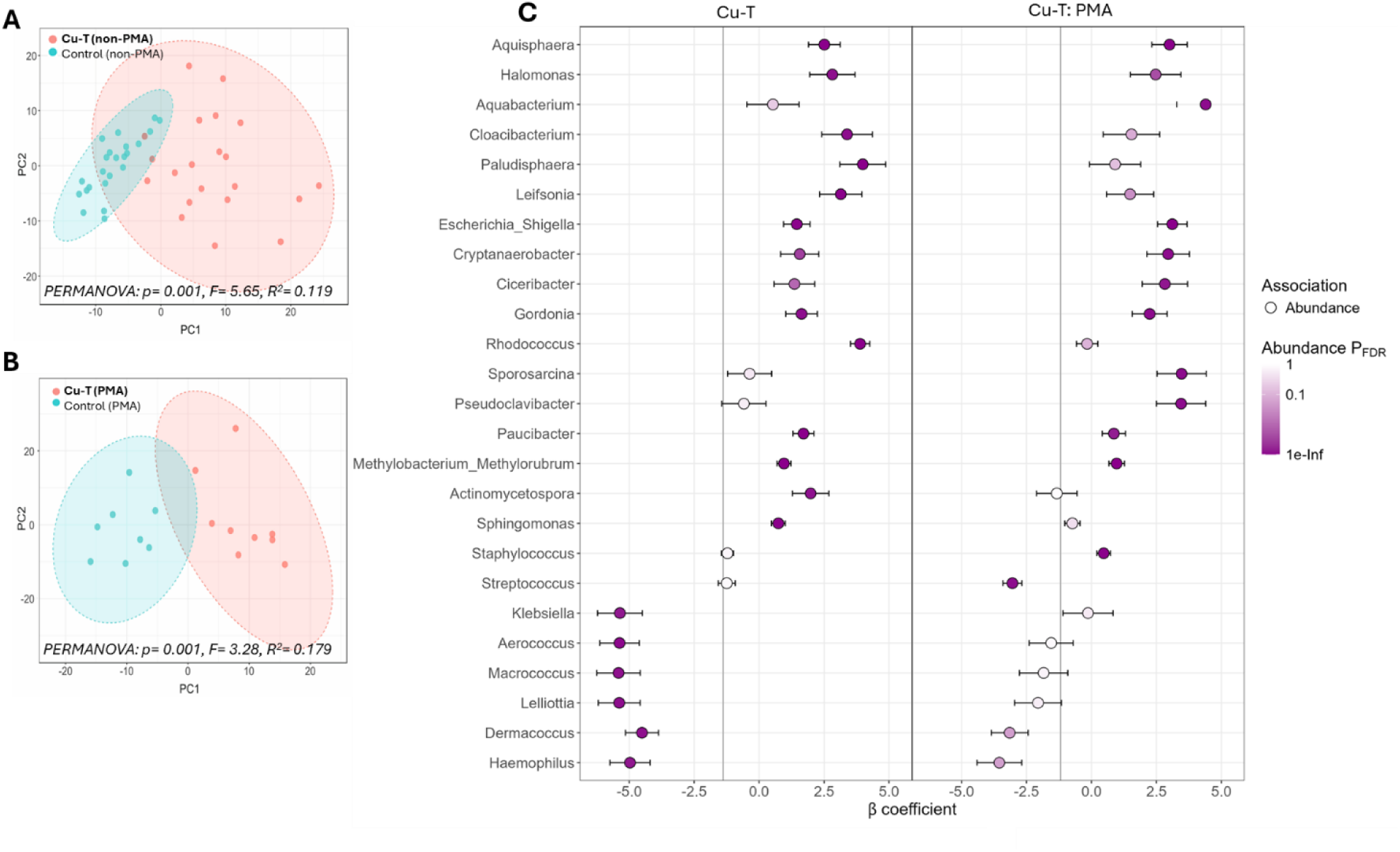
Beta diversity of non-PMA (A) and PMA-treated (B) bacterial communities on copper (Cu-T) and control basket handles, visualized by principal component analysis (PCA), with statistical significance assessed using permutational multivariate analysis of variance (PERMANOVA). Multivariable MaAsLin3 analysis (C) showing β coefficients, abundance effect sizes, and FDR-adjusted significance for bacterial genera associated with Cu-T and control surfaces, integrating results from both PMA-treated and non-PMA surface samples.

#### Community structure and genus-level associations

As shown above, basket handles onto which Cu-T was applied had the most varying bacterial community (Fig. 6) with *Staphylococcus* and *Streptococcus* being the most abundant genera. Similar to control basket handles the core genera on Cu-T coated basket handles involved *Streptococcus*, *Staphylococcus*, *Sphingomonas*, *Methylobacterium*, *Cutibacterium*, *Corynebacterium*, *Micrococcus*, and *Rhodococcus*, that were present at 100% prevalence on both surface types (Fig. S8).

Across both non-PMA and PMA treated surface samples, the microbial communities on control and Cu-T surfaces were dominated by *Staphylococcus*, although its relative abundance was consistently lower on Cu-T basket handles (Fig.7I). Other common genera including *Streptococcus, Cutibacterium, Corynebacterium, Paracoccus, Pseudomonas and Micrococcus* were present on both control and Cu-T surfaces but generally had lower relative abundances, with most showing small reductions on Cu-T. In contrast, *Rhodococcus* showed increased abundance on Cu-T surfaces as compared to control, suggesting enhanced tolerance or selective enrichment under copper exposure.

Multivariable MaAsLin3 results showed that microbial composition of basket handles was significantly affected by surface type (Cu-T versus control surfaces) and PMA treatment (FDR q < 0.10) (Fig. 8C). Cu-T surfaces were associated with 77 significant genera, including 39 enriched and 38 depleted compared to control surfaces. The genera that showed reduced abundances on Cu-T surfaces compared with control included *Macrococcus* (effect size (β) = −5.41), *Lelliottia* (−5.38), *Aerococcus* (−5.37), *Klebsiella* (−5.36), *Moraxella* (−5.19), *Psychrobacter* (−4.98), *Haemophilus* (−4.97), *Porphyromonas* (−4.95), *Abiotrophia* (−4.80), and *Brachybacterium* (−4.64). In contrast, Cu-T surfaces showed enrichment in *Paludisphaera* (3.99), *Rhodococcus* (3.89), *Cloacibacterium* (3.39), *Porphyrobacter* (3.18), *Leifsonia* (3.14), *Afipia* (2.95), *Halomonas* (2.82), *Aquisphaera* (2.51), *Aliterella* (2.24), *Rhodoblastus* (2.17), *Marisediminicola* (2.16), and *Limnobacter* (2.11) as compared to the control surfaces.

PMA treatment showed significant changes in abundance of 38 genera (26 enriched, 12 depleted) (Fig.8C). The genera most strongly enriched in PMA-treated surface samples included *Aquabacterium* (4.40), *Rubritepida* (3.59), *Sporosarcina* (3.47), *Pseudoclavibacter* (3.45), *Escherichia–Shigella* (3.11), *Aquisphaera* (3.01), *Cryptanaerobacter* (2.95), *Rummeliibacillus* (2.91), *Ciceribacter* (2.83), *Halomonas* (2.47), *Ilumatobacter* (2.27), and *Gordonia* (2.24), with additional increases for *Microbacterium*, *Tabrizicola*, *Ralstonia*, and *Leifsonia*. The genera that showed reduced abundances following PMA treatment included *Dietzia* (−5.20), *Prevotella* (−4.44), *Leuconostoc* (−4.31), *Noviherbaspirillum* (−3.89), *Haemophilus* (−3.55), *Brochothrix* (−3.23), *Dermacoccus* (−3.15), *Streptococcus* (−3.04), *Veillonella* (−2.84), and *Pseudomonas* (−2.75).

### Effect of TiO_2_-based coatings on microbiota of kindergarten tables

#### Microbial load and diversity

The presence of light-activated TiO_2_-C significantly lowered bacterial CFU/cm² on kindergarten tables compared with controls (p<0.05; Fig. 9A). The same trend was observed when 16S rRNA gene copies on TiO_2_-C and control surfaces (p<0.05; Fig. 9B) were compared. PMA treatment decreased the 16S rRNA copies on both, control tables in the kindergarten as well as on TiO_2_-C coated tables. When 16S rRNA gene copy number comparisons were made between PMA treated communities on control and TiO_2_-C surfaces a clear decrease of bacterial DNA on TiO_2_-C surfaces was detected (p<0.05; Fig. 9E).

**Figure 9.**
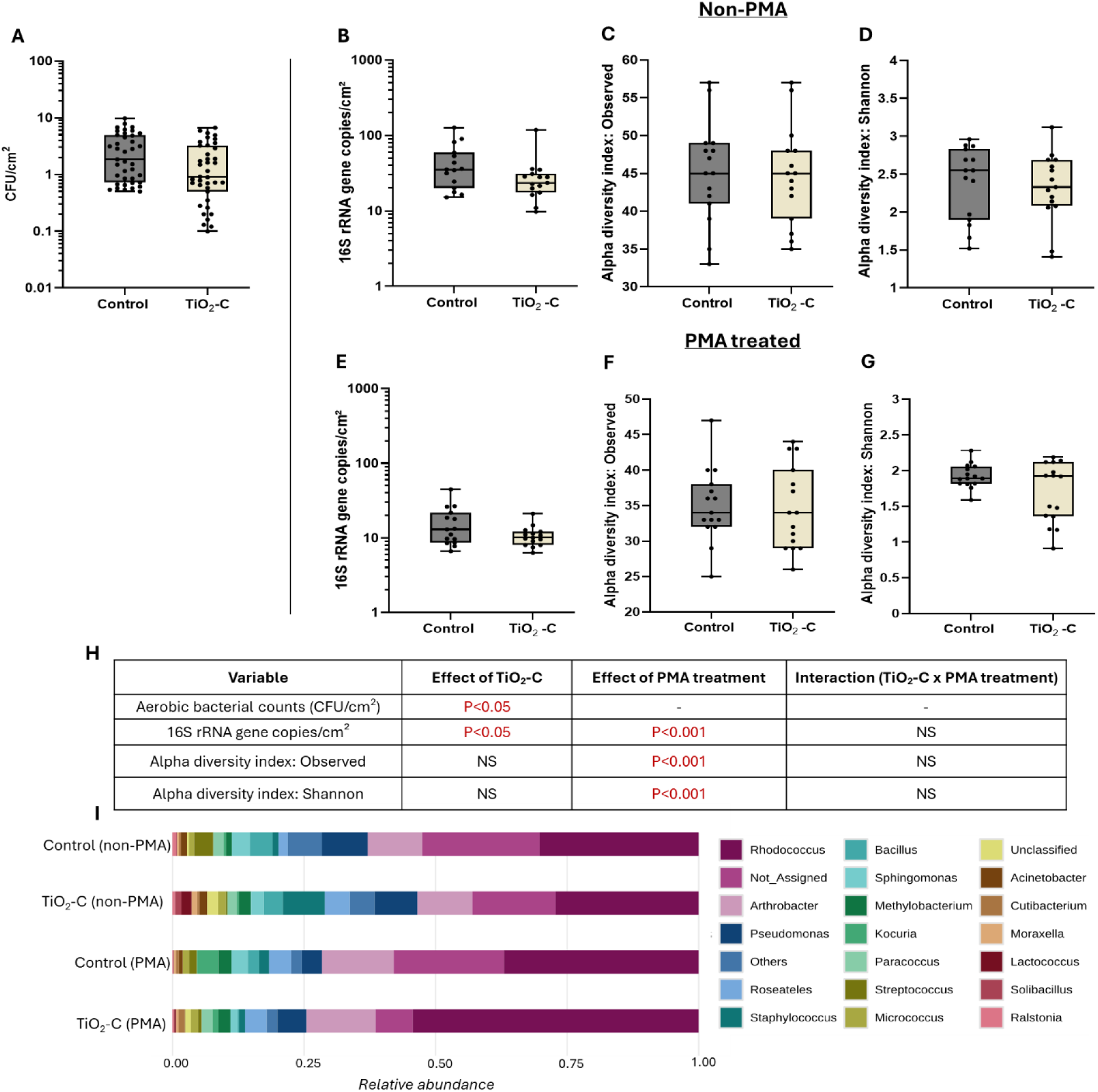
Comparison of bacterial load and community analysis on TiO_2_-based (TiO_2_-C) and control surfaces. Aerobic bacterial counts (CFU/cm²) (A), 16S rRNA gene copies per cm² for non-PMA (B) and PMA(E), observed ASV for non-PMA (C) and PMA(F), Shannon diversity index for non-PMA (D) and PMA (G) treated surface samples. Box-and-whisker plots show the median (center line), interquartile range (25th–75th percentiles; box), and minimum-maximum values. (H) Summary of statistical tests evaluating the effects of TiO_2_-C on CFU counts, 16S rRNA gene abundance, observed ASV, and Shannon diversity. One-way ANOVA was applied to CFU data, and two-way ANOVA was applied to Yeo–Johnson–transformed data to test the effects of TiO_2_-C, PMA treatment, and their interaction. NS = not significant (p ≥ 0.05). (I) Relative abundance of the top 20 genera across control and TiO_2_-C surfaces for PMA and non-PMA datasets; stacked bars represent merged proportional composition of bacterial genera from all surface samples per group with colors indicating individual genera. Relative abundance profiles of genera for individual surface samples are presented in Fig. S9.

Alpha diversity analysis showed no significant differences in community richness (Observed ASV; Fig.9C, 9F) or evenness (Shannon index; Fig.9D, 9G) between the TiO_2_-C and control surfaces in both PMA treated and non-PMA surface samples. This indicates that the overall diversity and structure of the bacterial communities was not affected by this TiO_2_-based surface coating.

Beta diversity analysis using PERMANOVA also demonstrated no statistically significant differences in community composition between the TiO_2_-C and control surfaces for both non-PMA (Fig. 10A) and PMA-treated surface samples (Fig. 10B). These overall results suggest that although TiO_2_-C may slightly reduce viable bacterial load, it does not substantially alter the overall microbial community structure under real-world conditions in the kindergarten environment.

**Figure 10.**
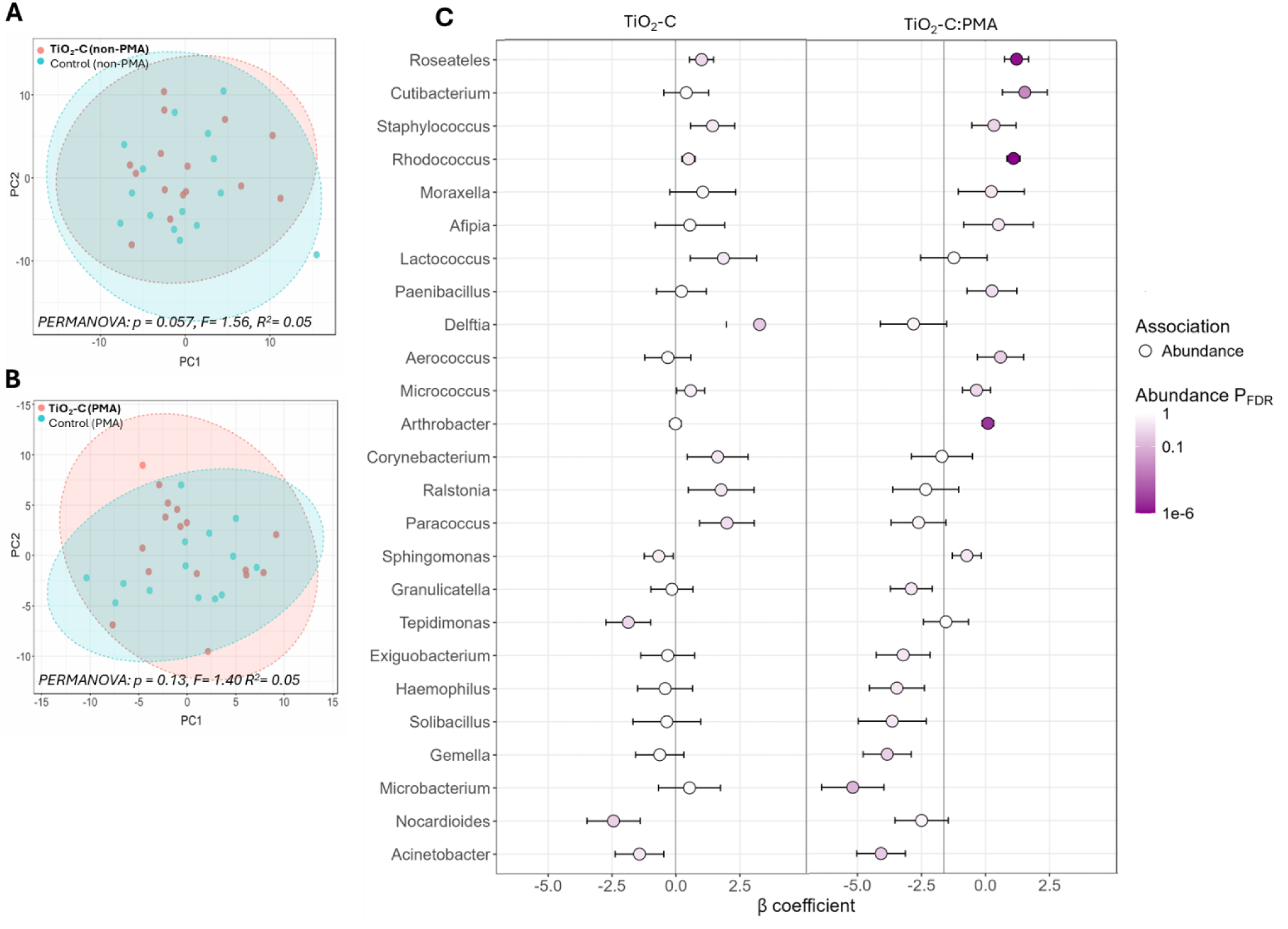
Beta diversity of non-PMA (A) and PMA-treated (B) bacterial communities from TiO_2_-based (TiO_2_-C) and control surfaces, visualized by principal component analysis (PCA), with statistical significance assessed using permutational multivariate analysis of variance (PERMANOVA). Multivariable MaAsLin3 analysis (C) showing β coefficients, abundance effect sizes, and FDR-adjusted significance for bacterial genera associated with TiO_2_-C and control surfaces, integrating results from both PMA-treated and non-PMA surface samples.

#### Community structure and genus-level associations

Core microbiome analysis revealed that the community compositions of the TiO_2_-C and control surfaces were broadly similar (Fig. S10). Across both PMA treated and non-PMA surface samples from TiO_2_-C and control, *Rhodococcus, Pseudomonas, Arthrobacter, Sphingomonas, Roseateles, Micrococcus, Methylobacterium,* and *Staphylococcus* were among the most consistently detected genera, indicating substantial overlap in both total and viable communities. The genus-level relative abundance patterns revealed that control and TiO_2_-C surfaces are dominated by *Rhodococcus, Arthrobacter,* and *Pseudomonas* (Fig.9I). Several genera including *Staphylococcus, Roseateles, Sphingomonas, Bacillus, Methylobacterium, Micrococcus, Streptococcus,* and *Cutibacterium* were present at lower abundances and showed only minor shifts between the control and TiO_2_-C. Multivariable MaAsLin3 analysis revealed PMA treatment as the only significant main effect influencing bacterial community composition (Fig. 10C). No significant associations were detected between TiO_2_-C surfaces and any bacterial genera, indicating no changes in genus-level relative abundances compared with control surfaces.

Comparison of PMA treated communities of kindergarten tables with non-PMA coated ones showed that in general, the survival of *Rhodococcus* (effect size (β) = 1.09), *Roseateles* (1.21), *Arthrobacter* (0.09), and *Cutibacterium* (1.53) on either control or TiO_2_-C-coated kindergarten tables was relatively high (Fig. 10C).

### Effect of silver-based surface on the microbiota of campus tables

#### Microbial load and diversity

Despite of the efficacy of Ag-based surface (Ag-S) in lab test showing the maximum of 2.4 log_10_ decrease of bacterial counts within 24 h (Fig. 2) no difference in microbial count collected from Ag-S surfaces in university campus compared with control surfaces was detected (Fig. 11A). Interestingly, 16S rRNA gene copy numbers were significantly lower on Ag-S than on control surfaces (p < 0.05; Fig. 11B) indicating clearly that the total aerobic counts measured with CFU counts only reflect a fraction of the actual surface microflora and even if CFU counts remain similar, the total bacterial count may have still changed. PMA treatment of surface communities decreased the 16S rRNA gene copies both, for control surfaces as well as for Ag-S surfaces. However, PMA treatment had a clearly stronger effect on reducing the 16S rRNA gene copies on control surfaces indicating that the relative presence of dead or damaged bacteria on those surfaces was higher. Due to this differential removal of bacterial cells from surfaces by PMA, the difference between PMA treated communities on control and Ag-S surfaces become non-existent (Fig. 11E).

**Figure 11.**
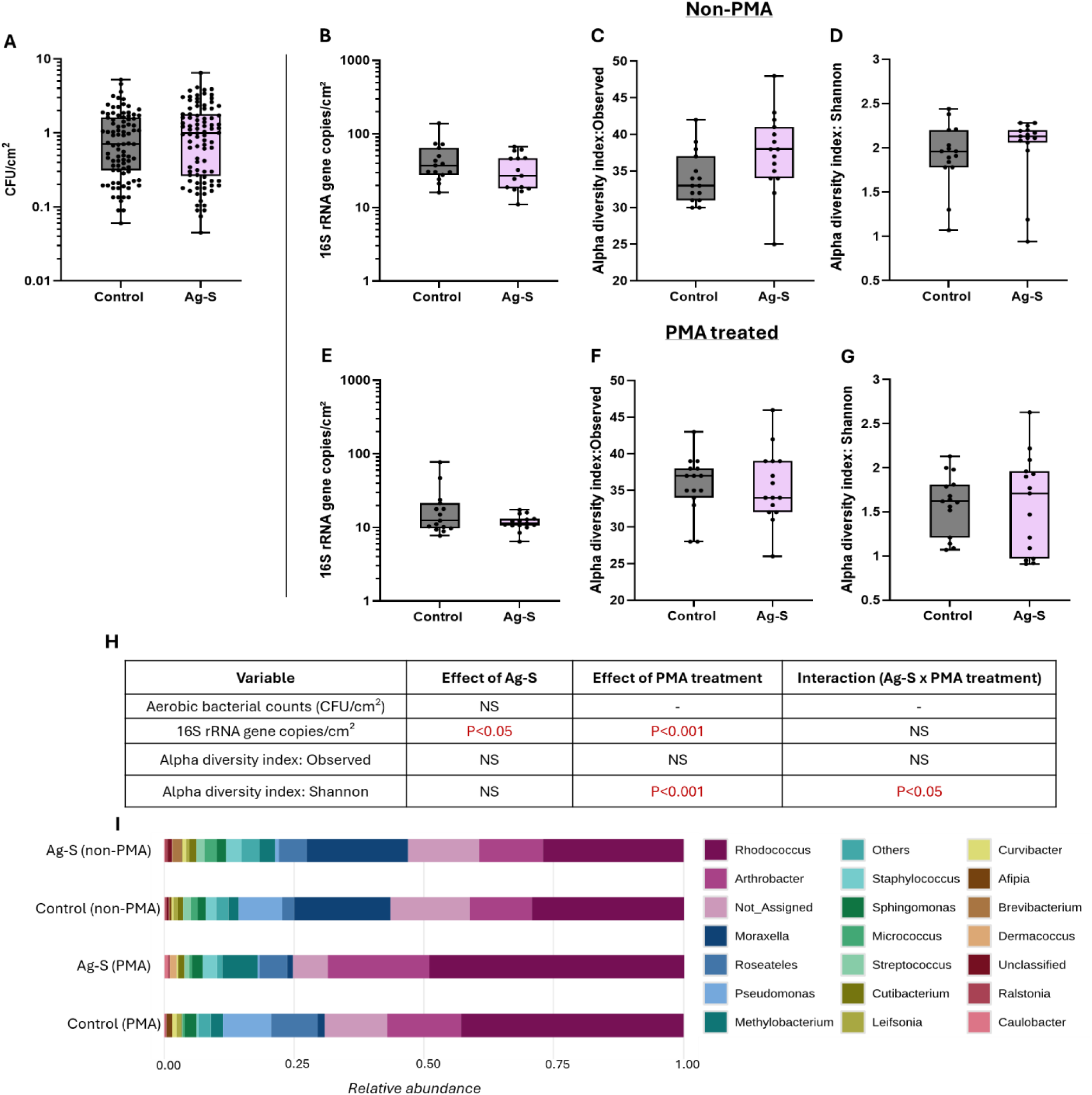
Comparison of bacterial load and community analysis on Ag-based (Ag-S) and control surfaces. Aerobic bacterial counts (CFU/cm²) (A), 16S rRNA gene copies per cm² for non-PMA(B) and PMA(E), Observed ASV for non-PMA(C) and PMA(F), Shannon diversity index for non-PMA(D) and PMA(G) treated surface samples. Box-and-whisker plots show the median (center line), interquartile range (25th–75th percentiles; box), and minimum–maximum values. (H) Summary of statistical tests evaluating the effects of Ag-S on CFU counts, 16S rRNA gene abundance, observed ASV, and Shannon diversity. One-way ANOVA was applied to CFU data, and two-way ANOVA was applied to Yeo–Johnson–transformed data to test the effects of Ag-S, PMA treatment, and their interaction. NS = not significant (p ≥ 0.05). (I) Relative abundance of the top 20 genera across control and Ag-S surfaces for PMA and non-PMA datasets; stacked bars represent merged proportional composition of bacterial genera from all surface samples per group with colors indicating individual genera. Relative abundance profiles of genera for individual surface samples are presented in Fig. S11.

Alpha diversity results based on the Observed ASV showed no statistically significant differences in the richness of genera between control and Ag-S surfaces. Compared with control, Ag-S had no effect on the Shannon diversity. PMA treatment resulted in a significant reduction in the Shannon diversity index compared with non-PMA samples (p < 0.001), demonstrating a clear distinction between total and viable microbial communities on both control and Ag-S surfaces. A modest interaction effect was also detected (p < 0.05) (Fig. 11H). As shown in Fig. 11D and 11G, PMA-treated control and Ag-S surfaces consistently exhibited lower Shannon diversity than their corresponding non-PMA samples, indicating subtle but measurable differences in the viable communities present on the both surfaces.

Beta diversity analysis showed no significant difference in non-PMA treated community composition between control and Ag-S surfaces (Fig.12A). However, PMA-treated communities on Ag-S and control surfaces exhibited a modest but significant separation between the two groups confirming the differences in the viable bacterial community (Fig.12B).

#### Community structure and genus-level associations

Core microbiome analysis revealed similar profiles between Ag-S and control tables at university campus, dominated by genera *Roseateles*, *Rhodococcus*, *Moraxella*, and *Arthrobacter* (Fig. S12). Other high-prevalence genera across both surface types included *Micrococcus*, *Staphylococcus*, *Methylobacterium*, *Sphingomonas*, *Cutibacterium*, and *Leifsonia*. The genus-level relative abundance patterns were also broadly similar between the Ag-S and control surfaces (Fig.11I) and the same taxa as in prevalence analysis appeared: *Rhodococcus, Moraxella, Arthrobacter*, *Pseudomonas* and *Roseateles*.

The multivariable MaAsLin3 analysis revealed that both PMA treatment and the presence of Ag-S surfaces had significant independent effects on the microbial communities found on university campus tables (Fig.12C). Compared with control surfaces, Ag-S surfaces showed a significant reduction in the relative abundance of *Actinomyces* (effect size, β = −2.37). Across both control and Ag-S surfaces, PMA treatment was associated with a substantially lower relative abundance of *Moraxella* (β = −3.46). In contrast, PMA-treated communities exhibited higher relative abundances of *Rhodococcus* (β = 0.94) and *Roseateles* (β = 1.33).

### Effect of SiQAC-based surfaces on the microbiota of cafeteria tables

#### Microbial load and diversity

Despite of the relative efficacy of SQ-C in lab tests (∼1 log_10_ CFU decrease in 1 h and maximum 2.4 log_10_ decrease in 24 h; Fig. 2) this coating did not cause any decrease in surface microbial load. On the contrary, aerobic bacterial counts (CFU/cm²) were significantly higher on SQ-C surfaces in cafeteria tables as compared to control tables (*p* < 0.05; Fig. 13A). The same trend was seen at 16S rRNA level: SQ-C increased the 16S rRNA gene copies on cafeteria tables (*p* < 0.001; Fig.13B). PMA treatment removed a fraction 16S rRNA genes from surface communities, but this removal was relatively even from both, control and SQ-C surfaces and therefore, also after PMA treatment SQ-C surfaces showed higher bacterial genome copies compared with control surfaces (*p* < 0.001; Fig. 13E).

**Figure 12.**
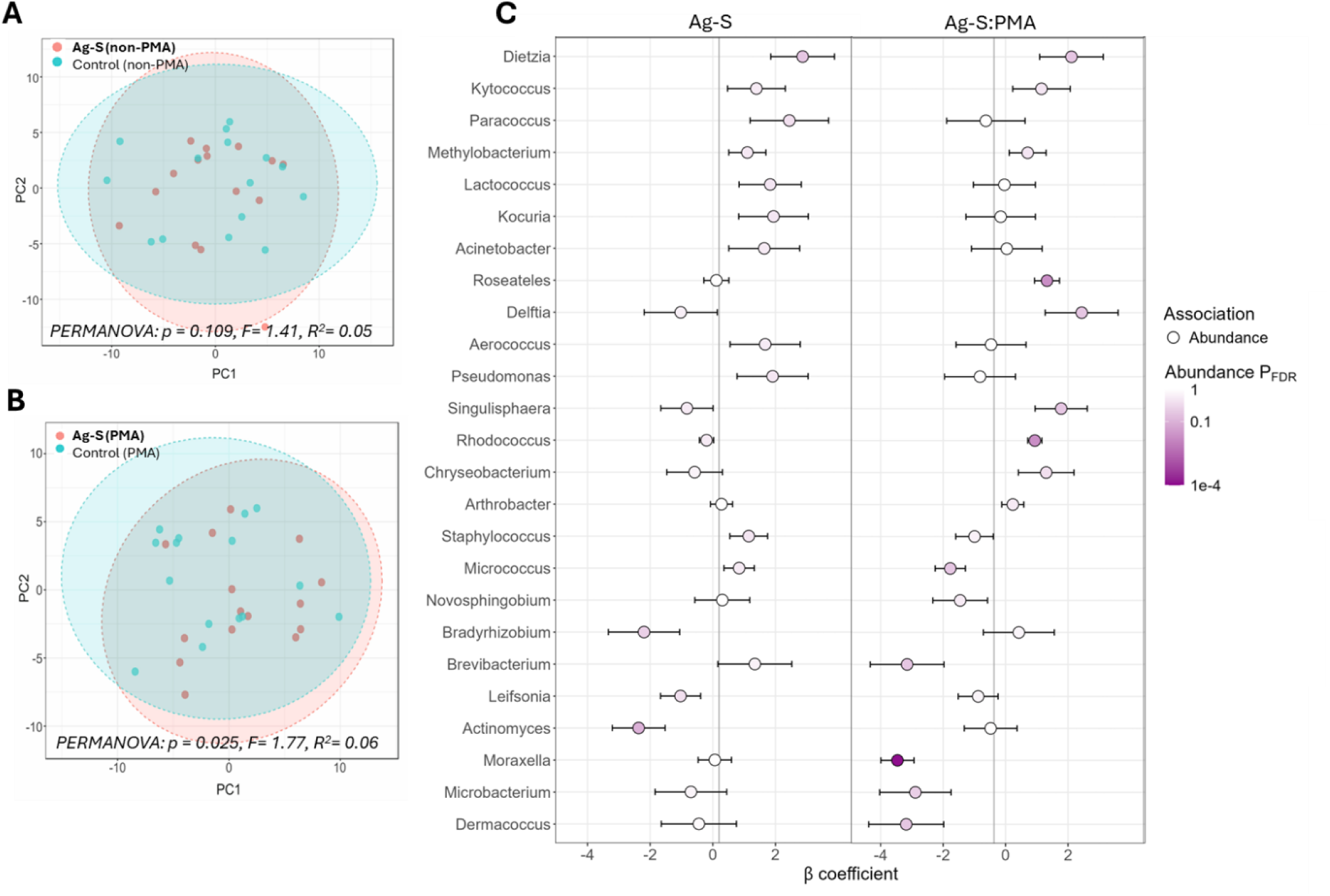
Beta diversity of non-PMA (A) or PMA-treated (B) bacterial communities from Ag-based (Ag-S) and control surfaces, visualized by principal component analysis (PCA), with statistical significance assessed using permutational multivariate analysis of variance (PERMANOVA). Multivariable MaAsLin3 analysis (C) showing β coefficients, abundance effect sizes, and FDR-adjusted significance for bacterial genera associated with Ag-S and control surfaces, integrating results from both PMA-treated and non-PMA surface samples.

**Figure 13.**
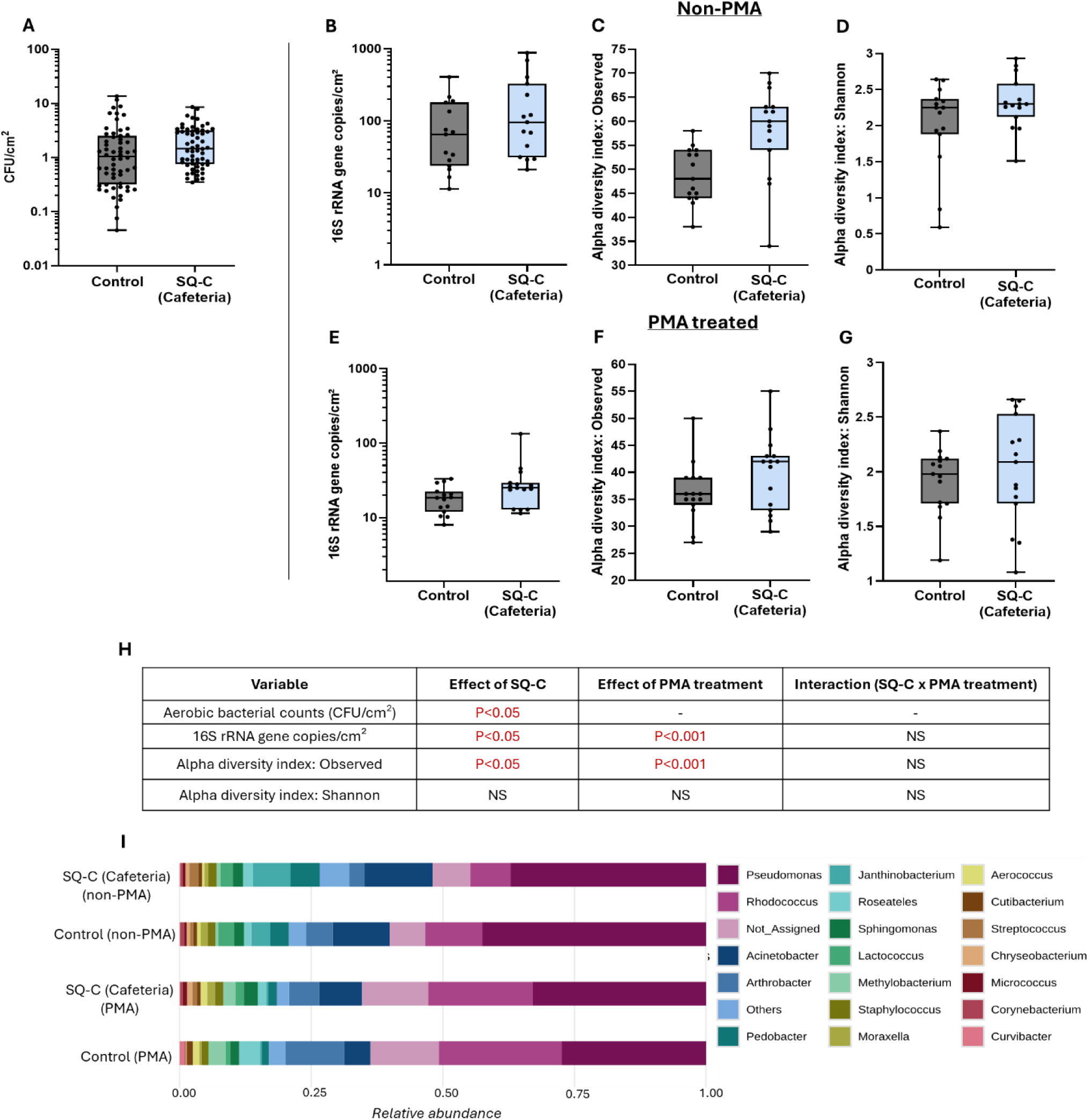
Comparison of bacterial load and community analysis on SiQAC based (SQ-C) and control surfaces in cafeteria. Aerobic bacterial counts (CFU/cm²) (A), 16S rRNA gene copies per cm² for non-PMA(B) and PMA(E), observed ASV for non-PMA (C) and PMA(F), Shannon diversity index for non-PMA(D) and PMA(G) treated surface samples. Box-and-whisker plots show the median (center line), interquartile range (25th–75th percentiles; box), and minimum–maximum values. (H) Summary of statistical tests evaluating the effects of SQ-C on CFU counts, 16S rRNA gene abundance, observed ASV, and Shannon diversity. One-way ANOVA was applied to CFU data, and two-way ANOVA was applied to Yeo–Johnson–transformed data to test the effects of SQ-C, PMA treatment, and their interaction. NS = not significant (p ≥ 0.05). (I) Relative abundance of the top 20 genera across control and SQ-C(cafeteria) surfaces for PMA and non-PMA datasets; stacked bars represent merged proportional composition of bacterial genera from all surface samples per group with colors indicating individual genera. Relative abundance profiles of genera for individual surface samples are presented in Fig. S13.

In line with the increased bacterial count, also alpha diversity analysis based on Observed ASV revealed significantly greater genus richness on SQ-C surfaces compared with control (Fig. 13 C and F) as indicated by two-way ANOVA analysis (Fig. 13H). Although the increase of Shannon diversity on SQ-C surfaces was not statistically significant (p > 0.05), the average on SQ-C was higher than on control surfaces (Fig. 13 D and G). Beta diversity analysis showed no significant difference in microbial community composition between non-PMA treated but also PMA-treated communities from control and SQ-C surfaces (Fig.14 A and B).

**Figure 14.**
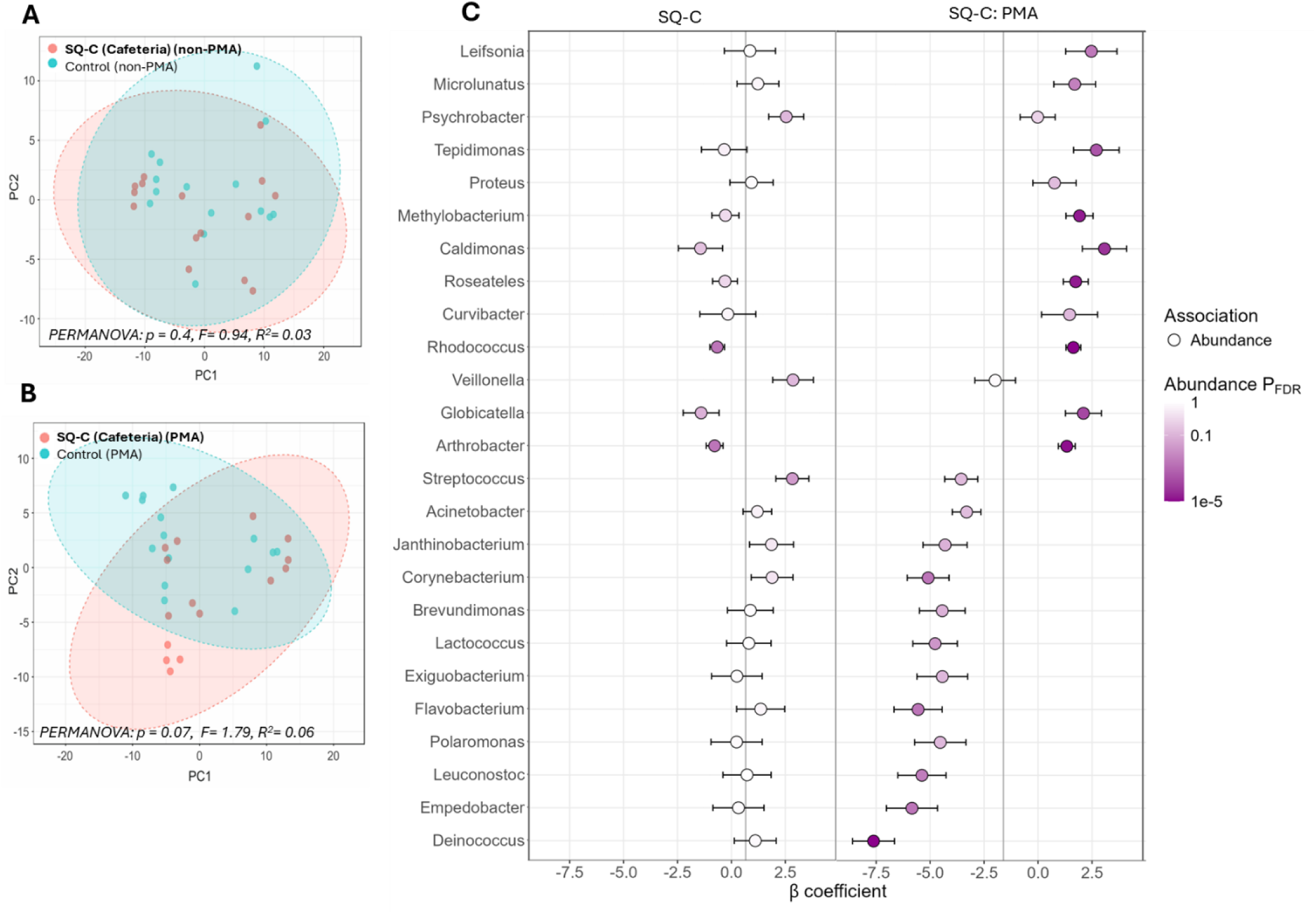
Beta diversity of bacterial communities on non-PMA (A) and PMA-treated (B) surface samples from SiQAC based (SQ-C) and control surfaces in cafeteria, visualized by principal component analysis (PCA), with statistical significance assessed using permutational multivariate analysis of variance (PERMANOVA). Multivariable MaAsLin3 analysis (C) showing β coefficients, abundance effect sizes, and FDR-adjusted significance for bacterial genera associated with SQ-C (cafeteria) and control surfaces, integrating results from both PMA-treated and non-PMA surface samples.

#### Community structure and genus-level associations

Core microbiome analysis revealed that SQ-C and control surfaces in cafeteria shared a broadly similar community structure, with most prevalent genera including *Pseudomonas, Rhodococcus, Arthrobacter, Staphylococcus,* and *Acinetobacter* (Fig. S14). *Pseudomonas*, followed by *Acinetobacter* and *Rhodococcus* were also the most abundant genera on both SQ-C and control surfaces (Fig. 13I), whereas *Arthrobacter* and *Roseateles* exhibited slightly higher relative abundances on control surfaces as compared to SQ-C surfaces. PERMANOVA analysis indicated that SQ-C surface did not fundamentally restructure the whole microbial community (Fig. 14 A and B). However, according to MaAsLin (Fig. 14C) specific taxa were significantly affected by the treatment. SQ-C coating was associated with higher relative abundance of *Streptococcus* (effect size (β) = 2.83) and lower abundances of *Globicatella* (−1.41), *Arthrobacter* (−0.78), and *Rhodococcus* (−0.66) as compared to control surfaces. PMA treatment of microbial communities collected from cafeteria surfaces was significantly associated with the reduction of *Deinococcus* (−7.62), *Empedobacter* (−5.84), *Flavobacterium* (−5.55), *Leuconostoc* (−5.38), *Corynebacterium* (−5.09) and *Lactococcus* (−4.77) both with increasing abundance of *Caldimonas* (effect size (β) = 3.08), *Tepidimonas* (2.70), *Leifsonia* (2.47), *Globicatella* (2.10), *Methylobacterium* (1.92) and *Rhodococcus* (1.64).

### Effect of SiQAC-based surfaces on microbiota of tables in animal clinic

#### Microbial load and diversity

On surfaces of animal clinic, the aerobic bacterial counts were consistently low with values <1 CFU/cm² (Fig.15A). However, no differences in CFU counts or 16S rRNA gene copy numbers were observed between control and SQ-C surfaces (Fig. 15 A and B). Differently from other study sites, PMA treatment did not cause any changes in the amount of bacterial 16S rRNA copies in surface samples from animal clinic (Fig. 15E). This was rather surprising, however could be an issue of detection limits as the number of surface-collected bacteria was low. Alpha diversity metrics based on Observed ASV and Shannon index showed similar richness (Fig. 15C, 15F) and evenness (Fig. 15D, 15G) of genera on control and SQ-C surfaces. The results of two-way ANOVA indicated that SQ-C treatment did not measurably influence overall bacterial load and the diversity (Fig. 15H).

**Figure 15.**
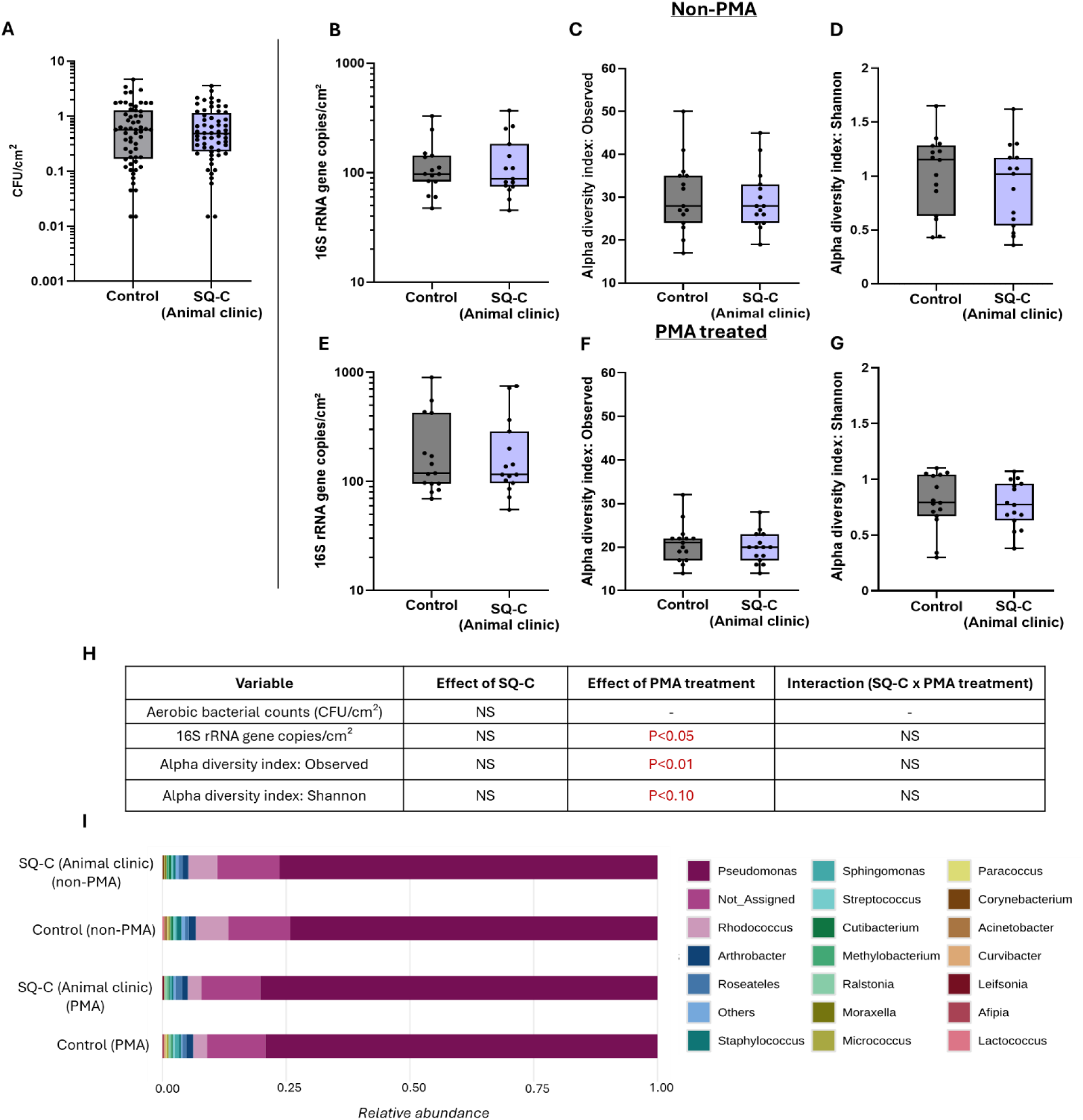
Comparison of bacterial load and community analysis on SiQAC based (SQ-C) and control surfaces in animal clinic. Aerobic bacterial counts (CFU/cm²) (A), 16S rRNA gene copies per cm² for non-PMA(B) and PMA(E), Observed ASV for non-PMA (C) and PMA(F), Shannon diversity index for non-PMA(D) and PMA(G) treated surface samples. (H) Summary of statistical tests evaluating the effects of SQ-C on CFU counts, 16S rRNA gene abundance, observed ASV, and Shannon diversity. One-way ANOVA was applied to CFU data, and two-way ANOVA was applied to Yeo–Johnson–transformed data to test the effects of SQ-C, PMA treatment, and their interaction. NS = not significant (p ≥ 0.05). Box-and-whisker plots show the median (center line), interquartile range (25th–75th percentiles; box), and minimum–maximum values. (I) Relative abundance of the top 20 genera across control and SQ-C (animal clinic) surfaces for PMA and non-PMA datasets; stacked bars represent merged proportional composition of bacterial genera from all surface samples per group with colors indicating individual genera. Relative abundance profiles of genera for individual surface samples are presented in Fig. S15.

Beta diversity analysis also revealed no differences in community structure associated with SQ-C surfaces in animal clinic compared to control surfaces (Fig. 16A, 16B). PERMANOVA results showed no significant separation between SQ-C and control surfaces in either non-PMA or PMA treated surface samples, further confirming that the SQ-C did not alter the overall microbial community composition.

**Figure 16.**
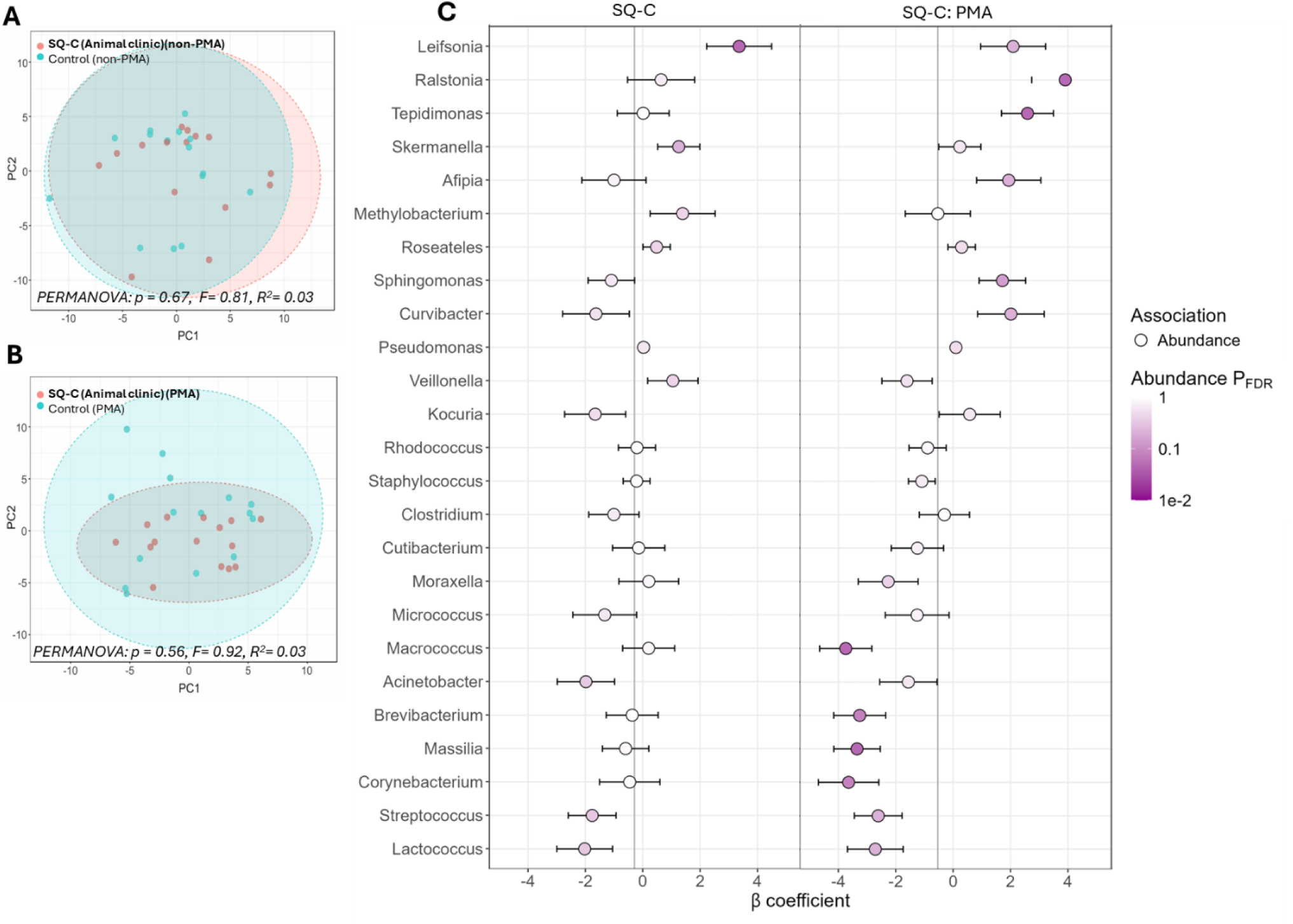
Beta diversity of bacterial communities on non-PMA (A) and PMA-treated (B) surface samples from SiQAC based (SQ-C) and control surfaces in animal clinic, visualized by principal component analysis (PCA), with statistical significance assessed using permutational multivariate analysis of variance (PERMANOVA). Multivariable MaAsLin3 analysis (C) showing β coefficients, abundance effect sizes, and FDR-adjusted significance for bacterial genera associated with SQ-C (animal clinic) and control surfaces, integrating results from both PMA-treated and non-PMA surface samples.

#### Community structure and genus-level associations

Core microbiome analysis revealed that both the SQ-C and control surfaces harboured a similar set of dominant taxa with *Pseudomonas, Rhodococcus, Arthrobacter*, and *Roseateles* being the most prevalent (Fig. S16). Other prevalent genera included *Staphylococcus*, *Streptococcus*, *Cutibacterium*, and *Methylobacterium*. The genus-level relative abundance profiles were similar for SQ-C and control surfaces, with *Pseudomonas* being the most dominant genus (Fig. 15I). Other abundant genera included *Rhodococcus*, *Roseateles*, *Arthrobacter*, *Cutibacterium*, and *Staphylococcus*.

The results of the multivariable MaAsLin3 model results showed that in animal clinic, *Leifsonia* (effect size (β) = 3.36) had higher relative abundance on SQ-C surfaces as compared to control surfaces (Fig. 16C). PMA treatment of animal clinic surface communities decreased the abundance of *Macrococcus* (−3.74), *Massilia* (−3.35), *Brevibacterium* (−3.26), and *Corynebacterium* (−3.64) but increased the abundance of *Ralstonia* (3.91) and *Tepidimonas* (2.59) (Fig. 16C).

## Discussion

High-touch public surfaces harbour diverse microbial communities, the composition of which is formed based on their interactions with the natural environment and humans, and further shaped by targeted human activity, such as cleaning or any other type of surface treatment. Due to increasing understanding of the importance of the microbiome of the built environment on human health [54], it becomes increasingly critical to recognise the factors affecting the composition of the ambient microflora. As the members of indoor microbiome, microbes present on touch surfaces, may be related with the possibility to transmit infectious diseases, and antimicrobial treatments are often applied with a goal to eliminate the infection-causing microbial groups. However, the efficacy of such treatments is usually not determined in real-life and their actual effect on surface microbiome is only poorly known. Therefore, this paper focused on four commercially available antimicrobial coatings that were introduced to high-touch surfaces in five study sites with a goal to study their effects on surface bacterial load and microbiome composition.

### Microbial load and diversity in real-life study sites differs

The five study sites that were used to assess the real-life efficacy of the four selected antimicrobial surfaces showed slightly varying starting bacterial load and diversity. Basket handles in a hardware store showed significantly higher CFU and 16S rRNA copies than any other study site. The reason was likely frequent use of shopping baskets and their infrequent cleaning practice compared with other sites where rigorous hygiene protocols are well established. Together with higher bacterial load also the alpha diversity, an indication for taxonomical richness was higher on basket handles compared with any other study sites. Beta diversity, an indication of differences in microbial composition, was significantly different in all the study sites. However, the clear difference of basket handles from other study sites was observed again as bacterial taxa from basket handles formed a clear distinct cluster in PCA analysis. The taxonomical profile identified on basket handles most resembled skin microflora [55]: the most prevalent genera included *Pseudomonas, Staphylococcus, Sphingomonas, Streptococcus* and *Cutibacterium*. On other study sites, the most prevalent genera were of natural origin (e.g., *Rhodococcus, Arthrobacter* and *Pseudomonas*) with some taxa associated with human skin (*Staphylococcus, Cutibacterium, Micrococcus*) and mucosa (*Streptococcus, Corynebacterium*). It is possible that the clear separation of basket handle microbiome from other study sites also partly reflects a batch effect, which may be driven by the fact that samples from basket handles were processed and sequenced as a separate batch from other surface samples, although all the details of protocol were kept identical throughout the study. However, the clearly differing CFU and 16S rRNA gene copy counts suggest that basket handles truly differed from other sites. All the most abundant bacterial genera that were identified in any of the sites have been associated with high-touch surfaces in earlier studies [46,56] indicating that the surface-associated microbiome observed in this study is representative of typical built-environment microbial communities shaped by repeated human contact and environmental deposition. Some members of the most abundant taxa (*Rhodococcus*, Cutibacterium) were also identified as a part of empty swab background, however, as the abundance of those groups in empty swabs (Table S1) significantly differed from their abundance in surface samples, then the background contamination was considered to have minimal effect on the results.

### Real-life efficacy of antimicrobial surfaces differs from lab test results

While all the surfaces were sufficiently effective under lab-scale testing following standard ISO protocols, their effect in real-life was modest and in case of some surfaces even non-existent. Clearly decreased antibacterial effect of a variety of surface types in real-life relevant low humidity conditions compared with the most used liquid-based ISO 22196 protocol has been observed also in a series of earlier studies, e.g., by Michels et al. (silver-based surfaces) [57], Kaur et al. (copper, silver and quaternary ammonium based surfaces) [8] and Maitz et al. (zinc-based surfaces) [9]. The reason for such a discrepancy is most likely decreased contact between bacteria and surface-released antimicrobial compounds but also decreased metabolic activity in dry conditions [58]. Despite of the relatively low efficacy of antimicrobial surfaces in real-life settings, their ability to decrease surface microbial numbers has been clearly reported. Copper surfaces have been shown to decrease the count of MRSA and VRE in hospital settings [18], copper and silver based coating reduced aerobic colony counts on antimicrobially coated high-touch areas such as bed rails and bedside tables in an orthopedic ward [59] and decreased microbial count on quaternary ammonium surfaces has been demonstrated [15]. Also, in this study we evidenced some decrease in bacterial count during real-life testing of copper (Cu-T) and TiO_2_-based (TiO_2_-C) surfaces. However, no decrease of surface bacterial count was measured on silver-containing Ag-S or quaternary ammonium-based SQ-C surfaces. These results were confirmed both, by aerobic colony counts but also by quantitatively determining 16S rRNA gene copies by qPCR.

### Copper-based surfaces showed strong antimicrobial efficacy and microbial community shifts

Cu-T-covered surfaces showed the fastest and most significant effect on bacterial viability, both in lab tests as well as in real-life study. This result was not surprising as according to earlier reviews; copper has been the only treatment that has significantly affected microbial surface load in real-life screening [60]. Cu-T-covered basket handles exhibited 67.3% reduction in aerobic CFU counts compared with the control. Also, a clear reduction in 16S rRNA copies proved that bacterial load on Cu-T basket handles was lower than on control. In order to rule out the possibility that qPCR-measured 16S rRNA sequences on Cu-T surfaces originated from dead bacteria, PMA treatment, which is commonly used for dead-alive distinction in molecular studies, was applied. Although recent reports have doubted the accuracy of this treatment in live-dead analysis [61–63], we performed PMA treatment for all real-life samples and presented their comparisons. Our lab-scale pre-optimization (results not shown) carried out with a mixture of dead and alive bacterial population showed relative efficacy of PMA in removing the signal of DNA from dead bacteria from PCR results. Also, in most of the real-life samples PMA treatment showed a different microbial load or microbial community structure indicating that the DNA of at least some dead or damaged bacterial cells was removed by this treatment. On both, Cu-T and control basket handles PMA treatment decreased 16S rRNA gene copies and compared with control surfaces after PMA-treatment, the number of 16S rRNA gene copies in Cu-T surfaces after PMA-treatment was significantly lower.

Microbiome analysis revealed reduced alpha diversity and distinct beta diversity profiles, indicating that Cu-T surfaces not only suppress microbial abundance but also alter the community composition. Cu-T surfaces reduce abundances of a broad range of genera, including several taxa commonly linked to human skin, the respiratory tract, and opportunistic infections, such as *Macrococcus*, *Aerococcus*, *Klebsiella*, *Haemophilus*, *Moraxella*, and *Porphyromonas*. The pronounced depletion of human-associated taxa suggests that Cu-T surfaces may be particularly effective at reducing microbial signatures associated with frequent human contact, a key consideration in high-touch public environments. In contrast, Cu-T surfaces showed relative enrichment of several environmental and stress-tolerant genera, including *Paludisphaera, Rhodococcus, Leifsonia, Aquisphaera, Halomonas,* and *Limnobacter*. These taxa are commonly detected in soil, water, or biofilm-associated habitats and are characterised by metabolic versatility and tolerance to oxidative and desiccation stress. Similar selective shifts toward environmentally resilient microorganisms have been reported on antimicrobial and metal-rich surfaces, suggesting that copper imposes a strong selective pressure favour stress-adapted taxa [64,65].

As mentioned earlier, PMA treatment of communities collected from both, control and Cu-T surfaces affected the community composition. In general PMA treatment enriched the presence of genera such as *Aquabacterium, Sporosarcina, Escherichia, Shigella, Aquisphaera,* and *Gordonia*, indicating a greater likelihood of persistence as viable cells on basket handles. The dominance of stress-tolerant and environmental taxa after PMA treatment aligns with previous observations that only a subset of deposited microorganisms remain viable under dry, nutrient-limited surface conditions, as reported for cleanroom and spacecraft-associated environments [51]. PMA-associated reduction in abundance of genera including *Dietzia, Prevotella, Leuconostoc, Haemophilus, Streptococcus,* and *Veillonella* suggests that most frequently the genera that did not survive on either dry Cu-T or control surface were of human origin. *Prevotella, Haemophilus, Streptococcus* and *Veillonella* have been found on (partly anaerobic) human mucosal membranes and gut [66] and therefore, their reduced survival on dry environmental surfaces could be expected. Also, earlier studies have shown that the viability of those genera is rapidly lost on abiotic surfaces [67].

### TiO_2_-based surfaces showed light-dependent activity with limited effect in the field study

The light-activated TiO_2_-C surfaces containing intrinsically UVA-activable TiO_2_ nanoparticles showed moderate antibacterial activity in laboratory conditions (2.5 log₁₀ reduction of *E. coli* after 4h). Despite of this moderate antibacterial activity TiO_2_-C surface was shown to decrease the bacterial load also in real-life: on kindergarten tables both CFU as well as 16S rRNA gene copies indicated that the overall bacterial burden was decreased. This trend was observed both, without and with PMA treatment of the surface communities.

TiO_2_-C surfaces did not exert a selective pressure strong enough to alter bacterial community structure at the genus level. The absence of significant taxonomical changes in microbial community suggests that the observed decrease in microbial load due to TiO_2_-C is not related with any specific effect on particular bacterial taxa but involves a non-selective antimicrobial mechanism. Indeed, it is well known that upon light activation TiO_2_ nanomaterials produce reactive oxygen species that rather unspecifically affect organic molecules, including bacterial cells [68]. The significant main effects of PMA treatment revealed increased abundances of *Rhodococcus*, *Roseateles*, *Arthrobacter*, and *Cutibacterium*, indicating that these taxa contribute to the viable fraction of the surface microbiome. Both *Rhodococcus* and *Arthrobacter* have been shown to survive the dessication and chemical stressors and have been found in frequently cleaned indoor surfaces, dust or soil [11]. The increased abundance of the skin-associated genus *Cutibacterium* highlights the role of human contact in shaping the viable microbial communities detected on surfaces [13].

### Silver-based surfaces showed minimal effect on the microbial load and composition

Even if Ag-S surfaces achieved a maximum reduction in viable bacterial counts after 24 h in lab-test, Ag-S surfaces in university campus had no effect on the CFU-based microbial load. At the same time, 16S rRNA-based gene copy numbers were lower on Ag-S surfaces than on controls indicating that the total aerobic CFU is the most relevant measure of the total bacterial load on surfaces. Indeed, when we compared the surface-retrieved total aerobic CFU counts from different study sites with qPCR-based 16S rRNA gene copies, there was no significant correlation (spearman r = 0.1856; p>0.05). Being in agreement with minor effect of Ag-S on microbial surface load, also alpha diversity was similar between Ag-S and control surfaces both without and with PMA treatment. These observations are further supported by beta diversity and relative abundance analyses, which demonstrated highly similar genus-level community compositions on Ag-S and control surfaces. The only identified difference concerned reduced abundance of *Actinomyces*, a genus commonly associated with human and animal mucosal microbiota [69] but also detected on a variety of environmental surfaces, on Ag-S surface. We may therefore hypothesize that Ag-S surfaces may more efficiently affect Gram-positive taxa [70], including genera such as *Actinomyces*.

Our finding demonstrating the increased loss of *Moraxella* from PMA treated surface communities suggests that this primarily human-associated genus exhibits limited survival outside the host and under stressful surface conditions typical of built environments as has been shown before [13, 14] Similar to kindergarten tables, including those with TiO_2_-C, *Rhodococcus* and *Roseateles* showed also higher survival on university campus tables. *Rhodococcus* is well known for its exceptional environmental resilience, which is supported by a robust, lipid-rich cell envelope containing mycolic acids, as well as high tolerance to desiccation, oxidative stress, and chemical exposure, which likely enhance membrane integrity and protect genomic DNA, resulting in preferential detection following PMA treatment [71].

### SiQAC coating had a neutral or even positive effect on bacterial surface load and community richness but highlighted some effect on community structure on cafeteria and animal clinic surfaces

Quaternary ammonium-based SQ-C was another example of a surface that showed efficacy in lab tests but not in real-life settings. Interestingly, the bacterial load, both in terms of CFU and 16S rRNA copies, increased on SQ-C coated cafeteria tables compared with control. SQ-C surfaces on cafeteria tables also showed higher microbial community richness compared to control. This increase in bacterial surface load and richness was surprising and unfortunately cannot be well explained. When SQ-C was applied to tables in the animal clinic, practically no change in surface bacterial load or community richness was observed. However, as the bacterial load on animal clinic samples was very low, it is possible that the differences could not be revealed because of the detection limits. Although higher CFU counts and increased richness were observed on SQ-C surfaces in the cafeteria, these changes were not accompanied by significant shifts in beta diversity, indicating that SQ-C did not fundamentally restructure the surface microbial community but rather influenced the relative representation of specific taxa within an otherwise stable community framework.

Compared with control surfaces, the SQ-C surfaces in cafeteria presented genus-specific shifts, with higher *Streptococcus* and lower *Globicatella, Arthrobacter,* and *Rhodococcus* abundances. In cafeteria, *Streptococcus* is plausibly linked to human occupancy and repeated re-introduction via hand contact and respiratory/oral shedding, which is consistent with built-environment studies showing strong occupant contributions on high-touch surfaces [72,47]. In contrast, the reduced abundance of *Arthrobacter* and *Rhodococcus* suggests that SQ-C may impose selective pressure even against stress-tolerant Actinobacteria, including taxa with documented desiccation tolerance [73]. In animal clinic, *Leifsonia* was more abundant on SQ-C surfaces, likely reflecting environmental introduction via soil and dirt carried by animals rather than a host-associated origin.

Similar to other study sites PMA treatment changed the abundance of some taxa on cafeteria and animal clinic surfaces. *Corynebacterium* and *Leuconostoc* that were less abundant in PMA-treated cafeteria communities, are human-related genera and likely do not preserve well on dry environmental samples. The same was true for moist environment and food-related *Lactococcus, Flavobacterium* and *Empedobacter* [74]. The genera that were more abundant in PMA treated cafeteria communities belonged to genera *Methylobacterium, Roseateles, Tepidimonas, Caldimonas, Rhodococcus, Microlunatus*, which are abundant in cleaning water, sinks, plumbing, damp residues and therefore, are more likely to survive harsh environmental conditions [75].

In animal clinic the taxa with reduced abundancy after PMA treatment included *Macrococcus*, *Massili*a, *Brevibacterium*, and *Corynebacterium*, which have been often associated with animal and human skin environment. *Corynebacterium* species are well-established members of the skin microbiota of both humans and animals, often dominating moist body sites and acting as commensals under normal conditions; however, some species have been implicated in opportunistic infections [76,77]. Similarly, *Macrococcus* has been isolated from the skin of animals and is frequently detected in comparative microbiome studies of equine and other mammalian skin surfaces, suggesting a consistent presence on epidermal habitats, although rarely causing disease [77]. *Massilia* is a gram-negative environmental genus frequently reported from soil, water, air, and dust microbiomes, and has been identified as a component of core indoor dust communities as well as in natural environments and soil habitats [78,79]. Other identified taxa, such as *Brevibacterium*, are known inhabitants of skin and environmental microbiomes and have been linked to diverse habitats including soil, dairy products, and human skin, or acting as rare opportunistic pathogens in susceptible individuals [80,81]. *Ralstoni*a and *Tepidimon*as belonged to taxa more abundant in PMA treated samples and may reflect the viability of environmental opportunists capable of persisting on clinic surfaces. Members of *Ralstonia* have been isolated from water and biofilm communities, and certain species have been recognised as emerging opportunistic pathogens in immunocompromised human patients, indicating that they may be relevant in clinical settings where environmental and host exposures overlap [82]. *Tepidimonas* has been generally categorized as an environmental genus but has been occasionally detected in the gastrointestinal microbiome of pigs [83]. Altogether those results indicate that the relative survival of human-related genera on SQ-C surfaces was generally lower and that of environmental taxa higher.

## Conclusions

This multisite field study demonstrates that the real-world performance of antimicrobial surfaces is highly context dependent and cannot be reliably inferred from standardized laboratory tests alone without field test validation. By integrating culture-based methods, quantitative molecular assays, 16S rRNA gene sequencing, and PMA-based viability profiling, this study demonstrated that copper surfaces showed the strongest and most robust antimicrobial performance among the four tested surfaces. Copper surfaces not only significantly reduced aerobic CFU counts and viable bacterial DNA but also induced clear and reproducible shifts in microbial community composition. These shifts were characterised by depletion of many human-associated and opportunistic taxa and relative enrichment of environmentally derived, stress-tolerant genera, indicating strong selective pressure exerted by copper under dry, nutrient-limited surface conditions. In contrast, TiO_2_-based light-inducible surfaces exhibited moderate reduction in bacterial load without substantial restructuring of microbial communities in the kindergarten environment. While laboratory tests confirmed light-dependent antimicrobial activity, field data suggest that under real-use indoor lighting and frequent recontamination, TiO_2_-based coating primarily lowers overall surface biomass rather than selectively reshapes community composition. The viable bacterial fraction on these surfaces was dominated by stress-tolerant environmental taxa and skin-associated bacteria, reflecting persistent human input and environmental resilience. Silver-based surfaces showed limited effects on surface microbiomes under real-world conditions, despite meeting antibacterial efficacy thresholds in laboratory testing. The community composition and diversity were similar to those of control surfaces, indicating that silver-based surfaces in this context mainly affected bacterial abundance rather than imposed strong selective pressure. Quaternary ammonium-based surfaces showed antibacterial efficacy in lab tests but proved inefficient in real use. Those surfaces did not reduce bacterial load nor significantly alter the overall microbial community structure under real-world field conditions while still showing some effect on microbial community composition. Overall, these findings further support that among any available antimicrobial surfaces, copper exhibits the most pronounced effect also in real use. Although some of the tested surface coatings affected the composition of surface microbiome, no specific enrichment of clinically important genera was detected suggesting that the use of these four tested antimicrobial surfaces is not specifically enriching bacteria that may potentially cause health hazard.

## Acknowledgments

Authors acknowledge Dr. Merilin Rosenberg for scientific discussions, Jan Lepamaa and Valdo Pajumaa for providing the materials for testing.

## Authors’ contributions

HK carried out most of the experimental part and majority of the writing, RK designed and carried out veterinary clinic related work, analyzed data and wrote the corresponding text, KT participated in qPCR and sequencing library preparation, MT designed the sequencing library preparation experiment and revised the corresponding text, JT advised on sequencing experiment and data analysis, carried out part of the respective data analysis and revised the text, MK coated and characterized the surfaces and wrote the respective text, DD characterized surface contact angles and wrote the respective text, VK advised on surface preparation and revised the respective text, LM participated in real-life sampling, DNA extraction, qPCR and respective data analysis, MA supervised data collection from kindergarten and university campus, MK and JT analysed participated in sample collection from kindergarten, analysed the cell counts and performed data analysis, AS designed and supervised the surface coatings and sample collection in veterinary clinic, AI designed the whole study, supervised all the aspects, wrote and revised the text.

## Ethics declaration

Not applicable as the study was carried out using commercially available surface coatings and no personalized information was collected.

## Consent for publication

Not applicable

## Availability of data and material

Sequencing datasets generated during the current study are available in the ENA repository, with the accession number PRJEB106797.

## Materials and methods for physico-chemical characterisation of surface coatings

To characterise the surface coatings, the coating layer was characterised on stainless steel. A Nova NanoSEM 450 (FEI, USA) with a 3 kV electron acceleration voltage and a secondary electron detector was used to visualise the top view of TiO_2_-based (TiO_2_-C), SiQAC-based (SQ-C) or copper-based (Cu-T) covered stainless steel coupons or pieces of Ag-based (Ag-S) surface. To obtain the cross-sectional SEM image, the surfaces were cut with an electric die grinder (Dremel, USA) to make it fit on a 90° SEM stub. The side of the surface was then scratched with a scalpel.

For elemental mapping, XPS analysis of the coupons with antimicrobial surfaces was performed with a SCIENTA SES-100 spectrometer (Scienta, Sweden) using 200 eV pass energy and nonmonochromatic Mg K-alpha X–rays (incident energy 1253.6 eV) from X-ray source XRE2. The electron take-off angle was 90° and the pressure inside the analysis chamber was less than 10–9 Torr during the data collection. The raw data was processed via the CasaXPS software (version 2.3.26). Data processing involved the removal of X-ray satellites and peaks originating from K-beta X-rays. FTIR spectrometer (Bruker Vertex V70, Germany) equipped with an A513 reflection accessory set at 45° was used to non-destructively measure the IR reflection-absorption spectrum of the SQ-C on stainless steel and stainless steel without coating. The resolution was set to 4 cm^-1^, 32 scans were performed, and the spectra were then averaged across 8 surfaces.

The water contact angles of the coatings were measured to evaluate the wetting properties (surface hydrophobicity). Measurements were performed using a custom-built moving platform based on a Thorlabs DT12 (Newton, NJ, USA) dovetail translation stage. A 2 µL droplet of deionised water was deposited onto each coated surface using a micropipette. After 10 s of stabilisation, the water droplet profile was photographed with a Canon EOS 650d camera (Ota City, Tokyo, Japan) using a MP-E 65 mm f/2.8 1–5 × macro focus lens. Contact angles were determined from images captured from four water droplets on the same surface using ImageJ software (version 1.8.0_172; NIH, Madison, WI, USA) with a contact angle measurement plugin [1]. Samples with contact angle < 90° were considered hydrophilic, while those above 90° were considered hydrophobic.

---

**Figure S1.**
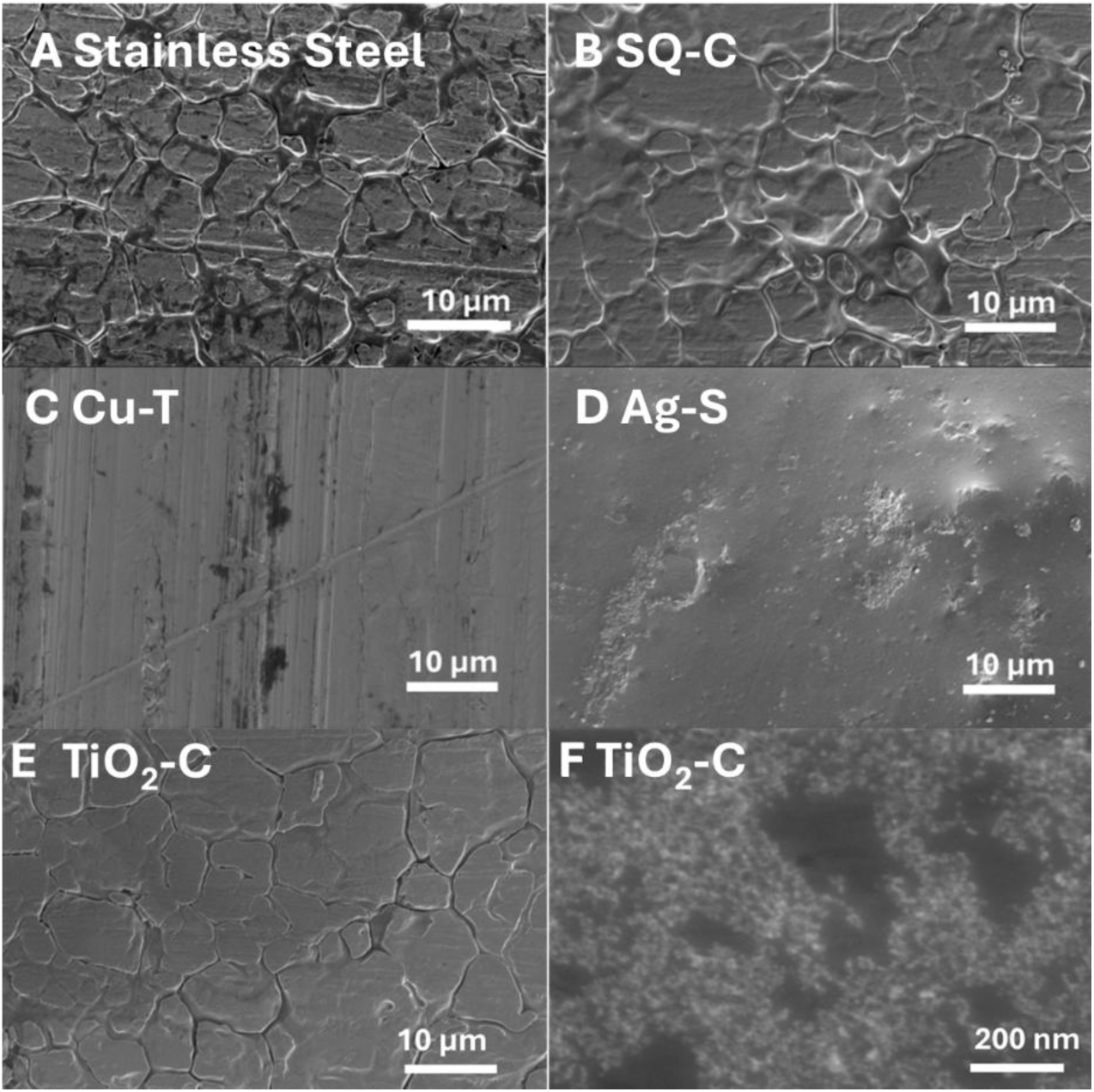
Scanning electron microscopy (SEM) images of surface coatings used in the study. At 50 000x magnification SiQAC-based (SQ-C) (B) and TiO_2_-based (TiO_2_-C) (E) coatings showed morphology that at large scale imitated the structure of stainless steel (A) but where from reduced contrast the presence of a thin but uniform coating could be assumed. A closer view of TiO_2_-C at 250 000x magnification (F) revealed the presence of 10-20 nm sized TiO_2_ nanoparticles on the surface of that coating. The appearance of the copper based (Cu-T) surface under (C) was in accordance with previously published SEM images of copper and the Ag-S surface (D) was visually smooth resembling polymer surfaces in general.

**Figure S2.**
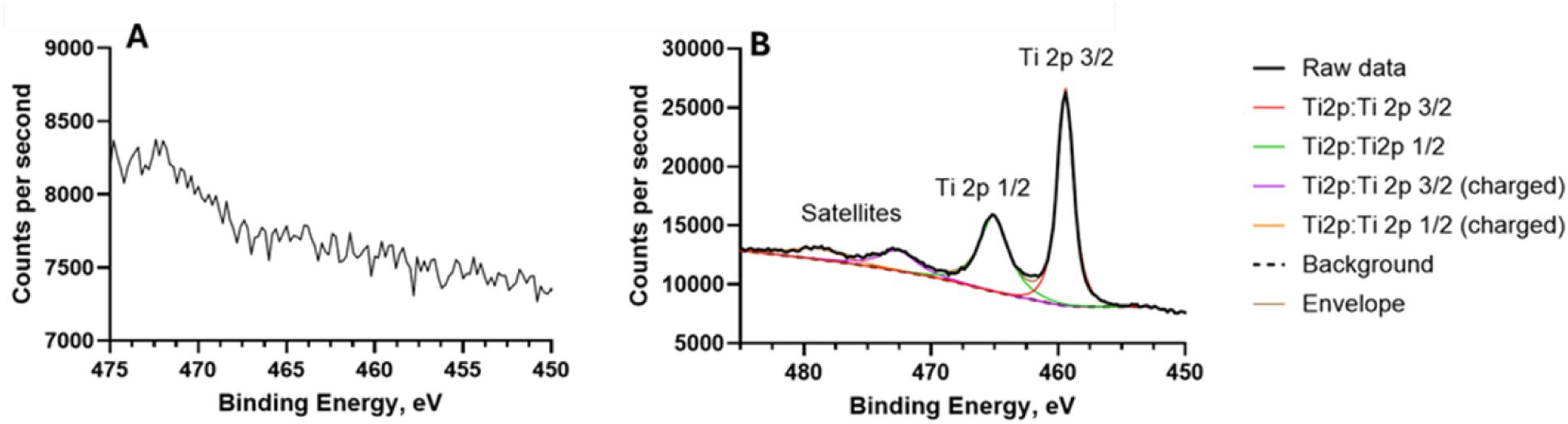
High-resolution Ti 2p X-ray photoelectron spectroscopy (XPS) spectra of (A) stainless steel and (B) the TiO_2_-based (TiO_2_-C) surface. The Ti 2p core-level spectrum consists of two spin–orbit components, Ti 2p₃/₂ and Ti 2p₁/₂, which arise from the splitting of the Ti 2p electron subshell due to total angular momentum coupling. In the high-resolution Ti 2p spectrum of the TiO_2_-C (Fig. S2B), a well-defined Ti 2p spin–orbit doublet is observed. The binding energy positions and splitting of these peaks are characteristic of Ti⁴⁺ species, confirming the presence of TiO₂ on the TiO_2_-C surface.

**Figure S3.**
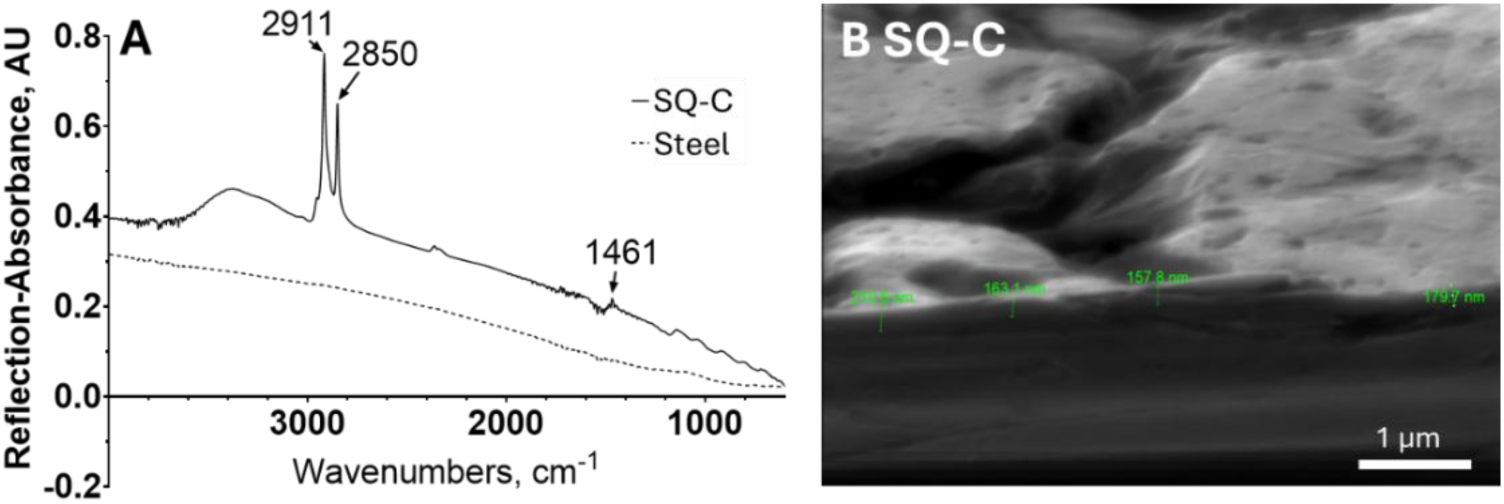
(A) Fourier transformed infrared (FT-IR) reflection absorption spectra of stainless steel and SiQAC-based (SQ-C) coating on stainless steel. The spectrum of the SQ-C exhibits characteristic absorption bands associated with quaternary ammonium compounds. The peak at 2911 cm⁻¹ is assigned to aliphatic –CH stretching, while the band at 1461 cm⁻¹ corresponds to C–N bond vibrations. These absorption features are consistent with quaternary ammonium compounds reported in the literature, confirming the presence of SQ-C on the stainless-steel surface. (B) Side-view SEM image of the SQ-C on stainless steel. Higher-magnification cross-sectional SEM images confirm the presence of a uniform, thin SiQAC layer. Based on the cross-sectional analysis (green numbers on B), the thickness of the SQ-C is estimated to be approximately 170–210 nm, as indicated by the scale bar.

**Figure S4.**
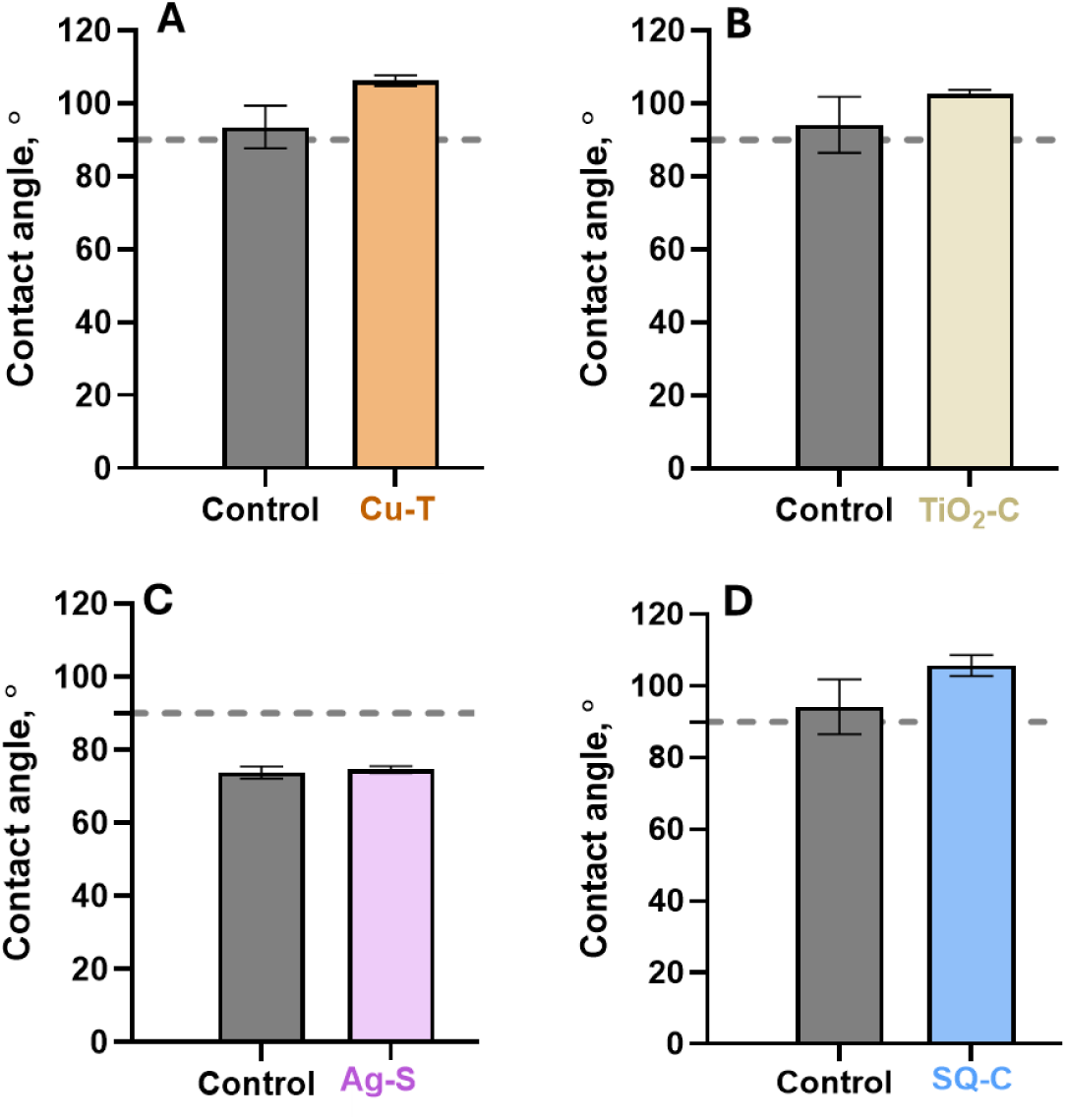
Contact angles of copper based (Cu-T) (A) TiO_2_-based (TiO_2_-C) (B), Ag-based (Ag-S) (C) and SiQAC-based (SQ-C) (D). Stainless steel was used as control for TiO_2_-C and SQ-C, plastic for Cu-T, and a control surface without Ag-additives was used as a control for Ag-S. Cu-T, TiO_2_-C and SQ-C surfaces exhibited water contact angles slightly above 90°, indicating a hydrophobic surface character (A, B, and D). In contrast, Ag-S and its corresponding control surface displayed contact angle values below 90°, demonstrating a hydrophilic nature (C). Although a slight increase in contact angle was observed for the Cu-T, TiO_2_-C,and SQ-C surfaces compared to control stainless steel, the overall differences were not statistically significant (p>0.05, t test with Welch’s correction). These results indicate that the antimicrobial surfaces do not drastically change the intrinsic surface hydrophilicity.

**Figure S5.**
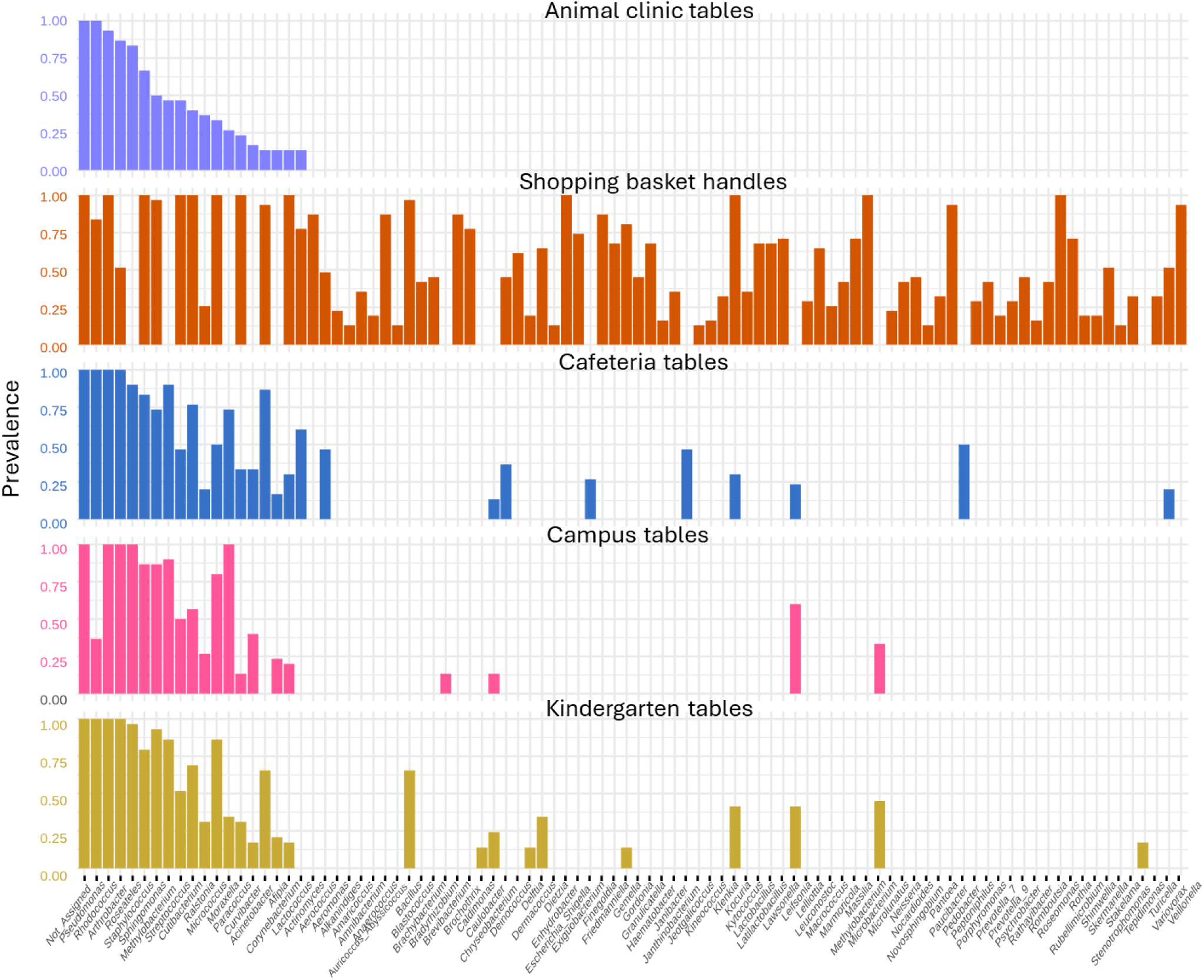
Prevalence of bacterial genera across samples collected from animal clinic tables, shopping basket handles, cafeteria tables, campus tables, and kindergarten tables. Bars represent the proportion of samples in which each genus was detected, applying a relative abundance threshold of 0.001 and a sample prevalence cutoff of 10%. Genera are displayed in descending order of prevalence within each condition.

**Figure S6.**
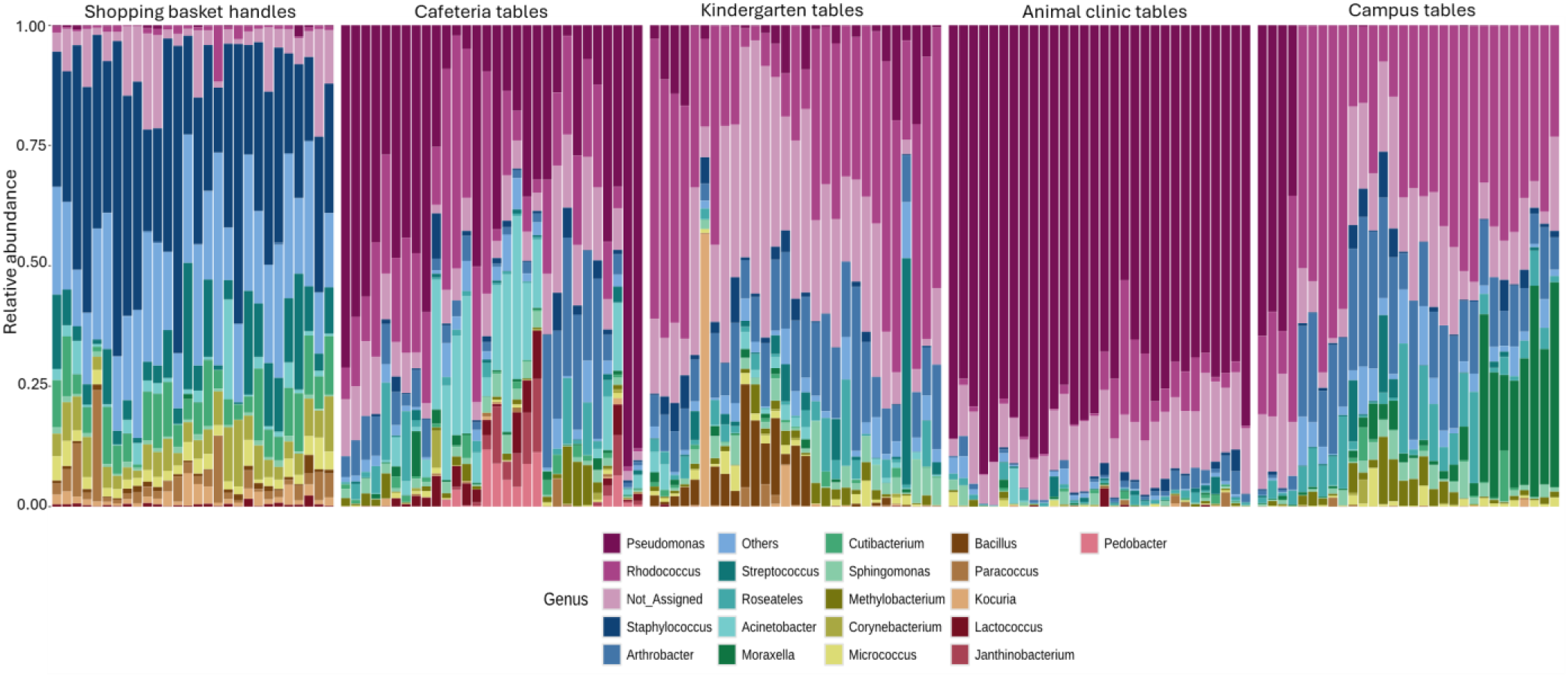
Relative abundance of the top 20 bacterial genera detected across control surface samples from shopping basket handles, cafeteria tables, kindergarten tables, animal clinic tables, and campus tables. Each stacked bar represents an individual surface sample, with bar height showing the relative abundance of each genus.

**Figure S7.**
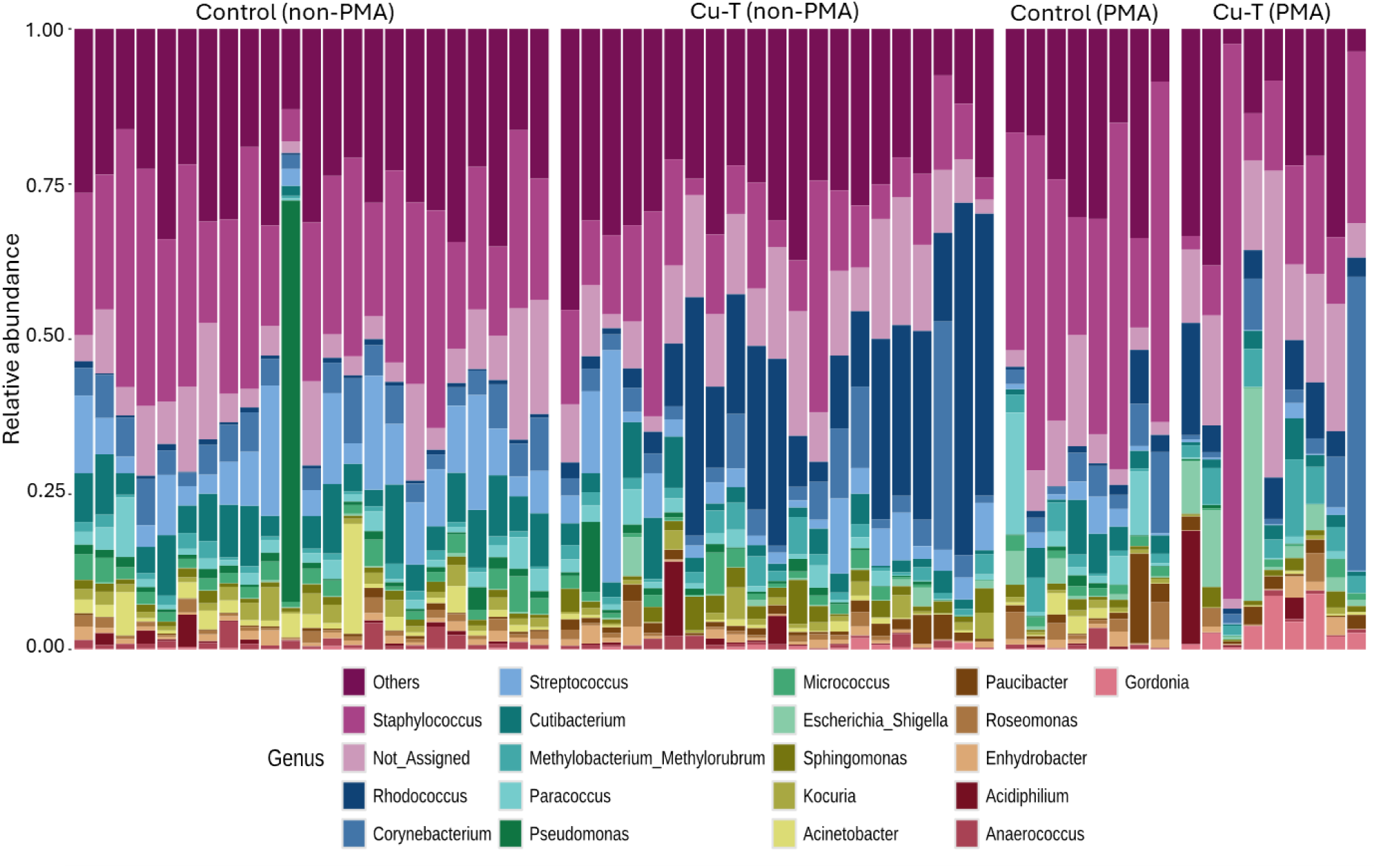
Relative abundance of the top 20 bacterial genera detected on shopping basket handles. Each stacked bar represents an individual surface sample on either non-PMA or PMA treated control or copper-based (Cu-T) surfaces, with bar height indicating the proportion of each genus within that sample.

**Figure S8.**
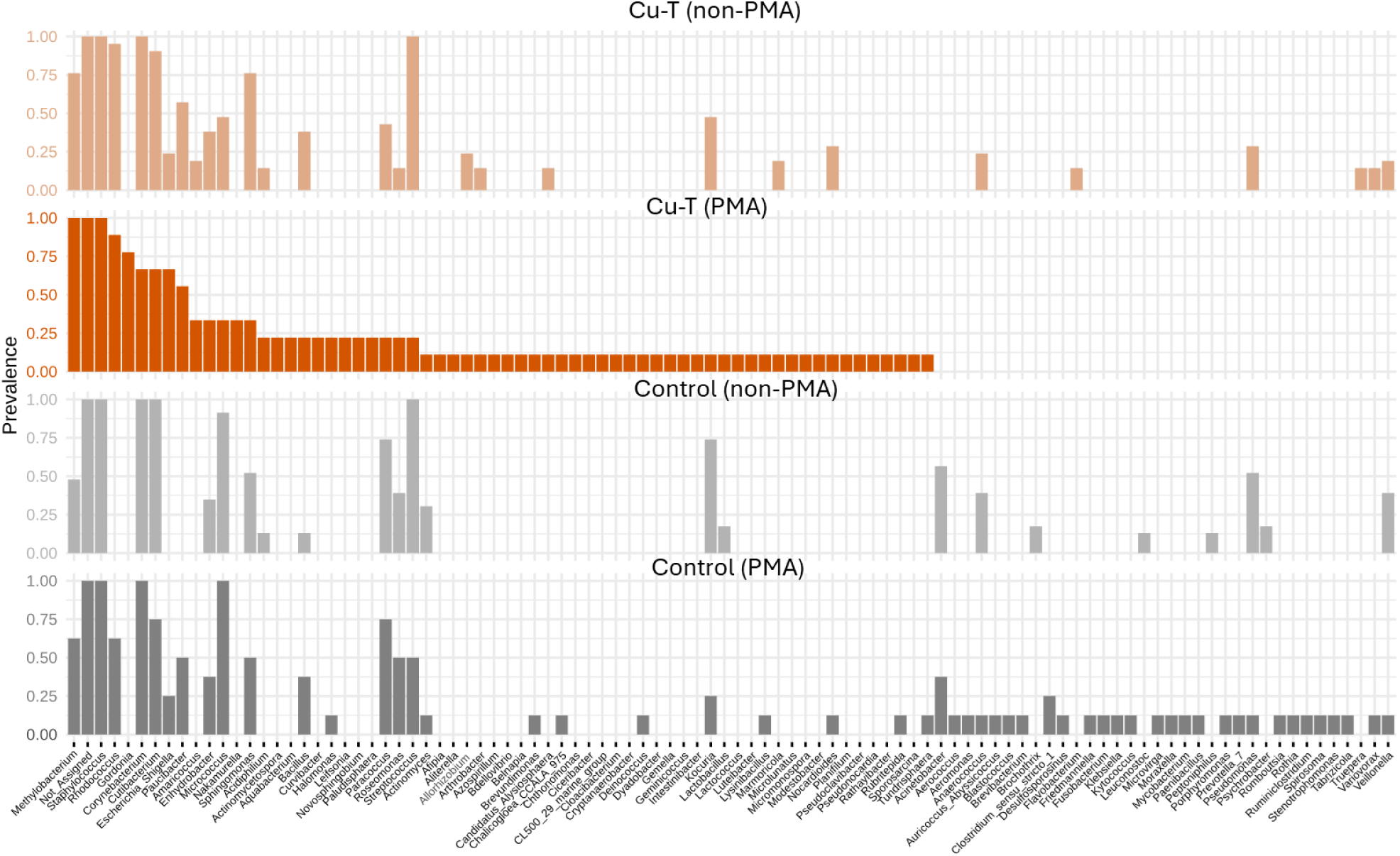
Prevalence of bacterial genera across samples collected from basket handles in the hardware store. Bars represent the proportion non-PMA or PMA treated control or copper (Cu-T) surface samples in which each genus was detected, applying a relative abundance threshold of 0.001 and a sample prevalence cutoff of 10%. Genera are displayed in descending order of prevalence within each condition.

**Figure S9.**
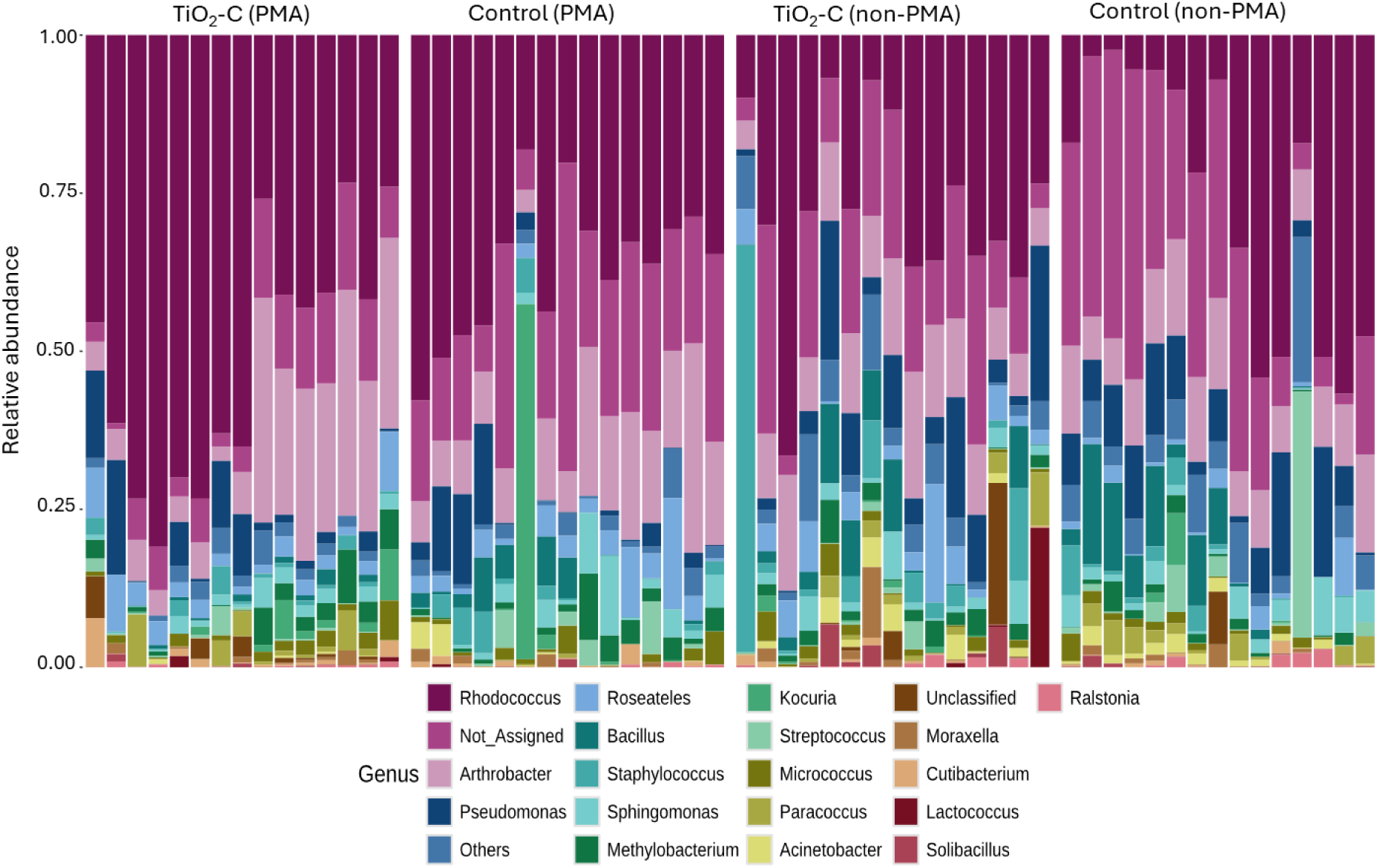
Relative abundance of the top 20 bacterial genera detected on kindergarten tables. Each stacked bar represents an individual surface sample on either non-PMA or PMA treated control or TiO_2_-based (TiO_2_-C) surface with bar height indicating the proportion of each genus within that sample.

**Figure S10.**
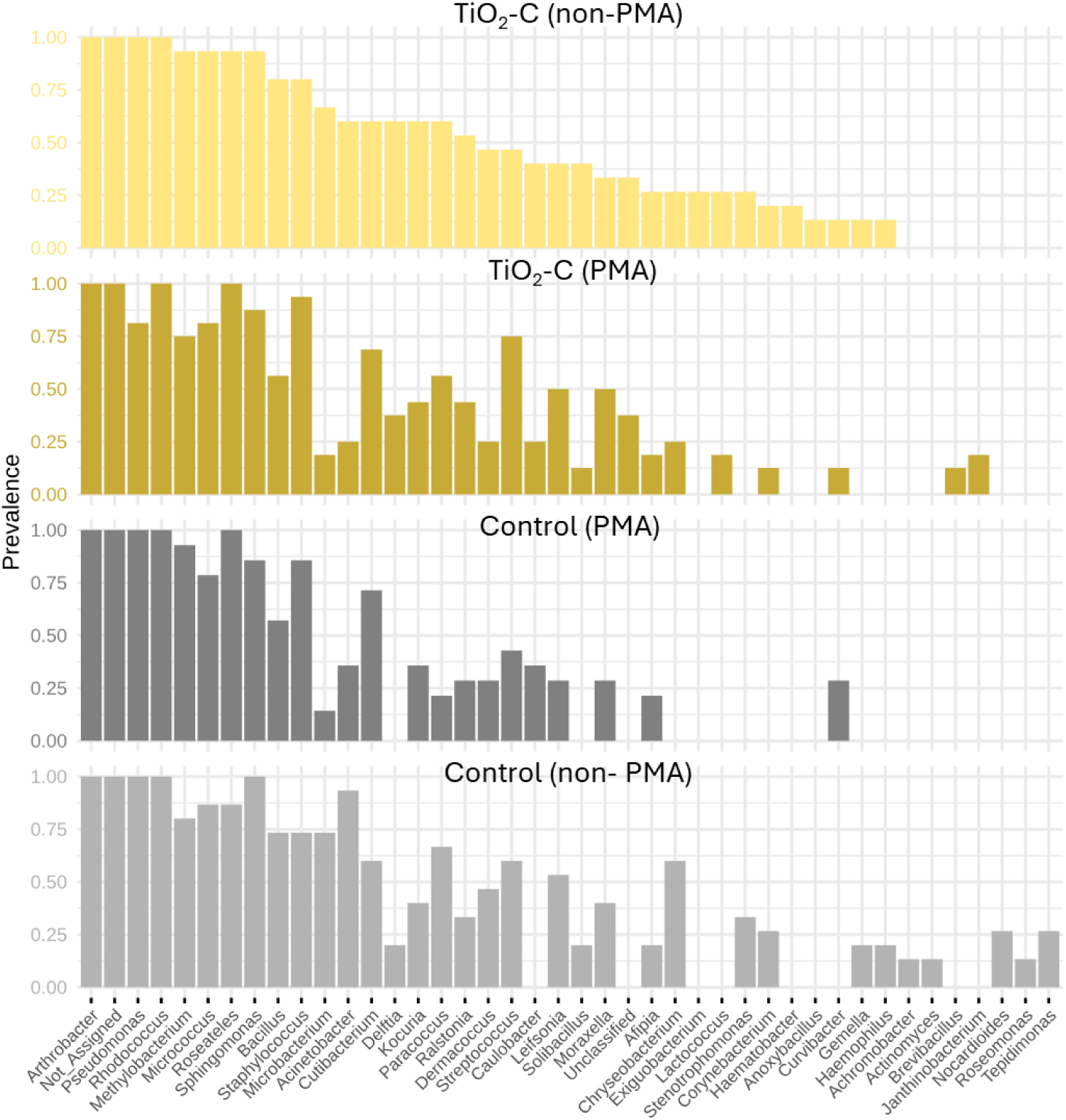
Prevalence of bacterial genera across samples collected from kindergarten tables. Bars represent the proportion either non-PMA or PMA treated control or TiO_2_-based (TiO_2_-C) surface samples in which each genus was detected, applying a relative abundance threshold of 0.001 and a sample prevalence cutoff of 10%. Genera are displayed in descending order of prevalence within each condition.

**Figure S11.**
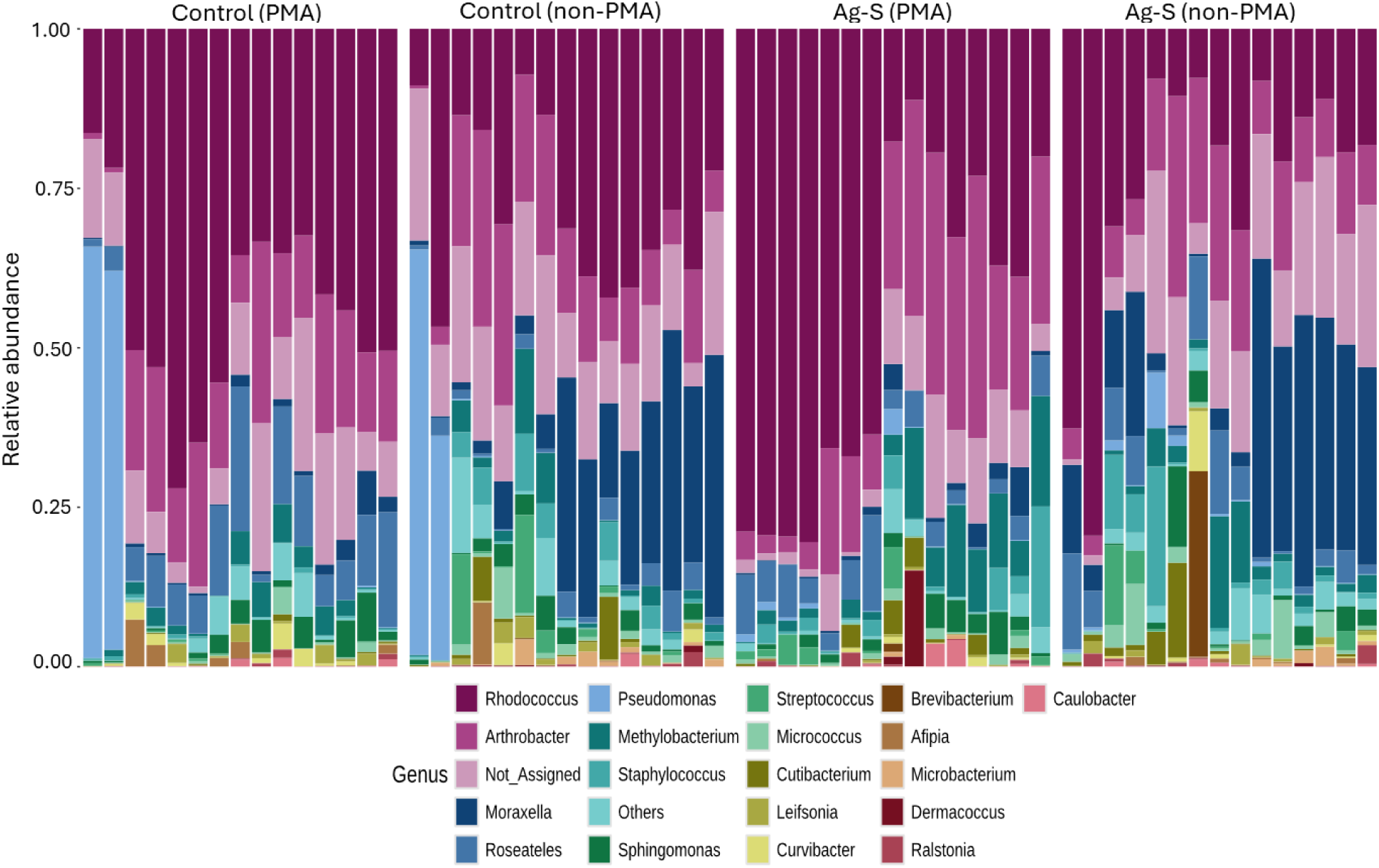
Relative abundance of the top 20 bacterial genera detected on tables in university campus. Each stacked bar represents an individual surface sample on either non-PMA or PMA treated control or Ag-based (Ag-S) surface with bar height indicating the proportion of each genus within that sample.

**Figure S12.**
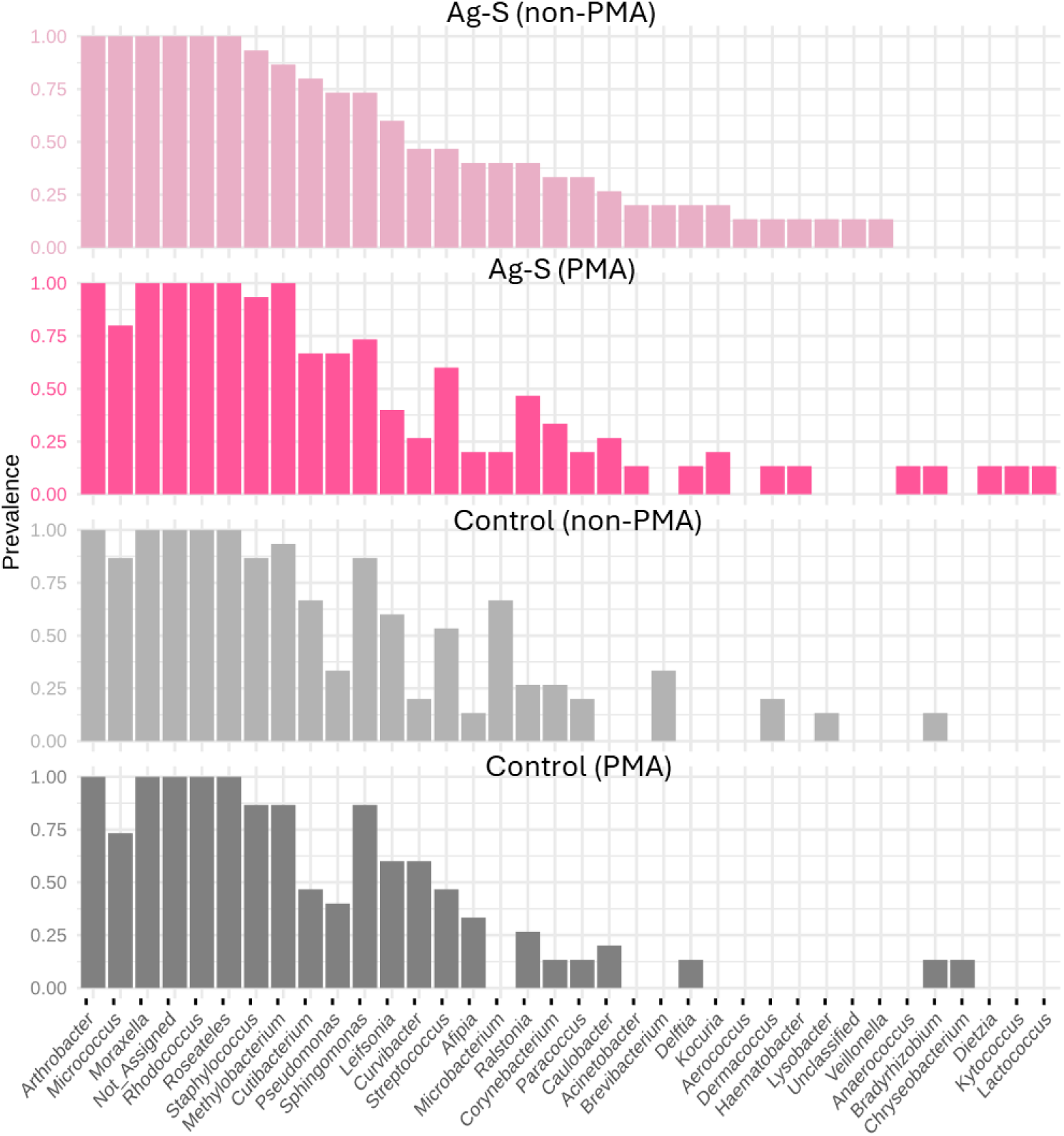
Prevalence of bacterial genera across samples collected from tables in university campus. Bars represent the proportion either non-PMA or PMA treated control or Ag-based (Ag-S) surface samples in which each genus was detected, applying a relative abundance threshold of 0.001 and a sample prevalence cutoff of 10%. Genera are displayed in descending order of prevalence within each condition.

**Figure S13.**
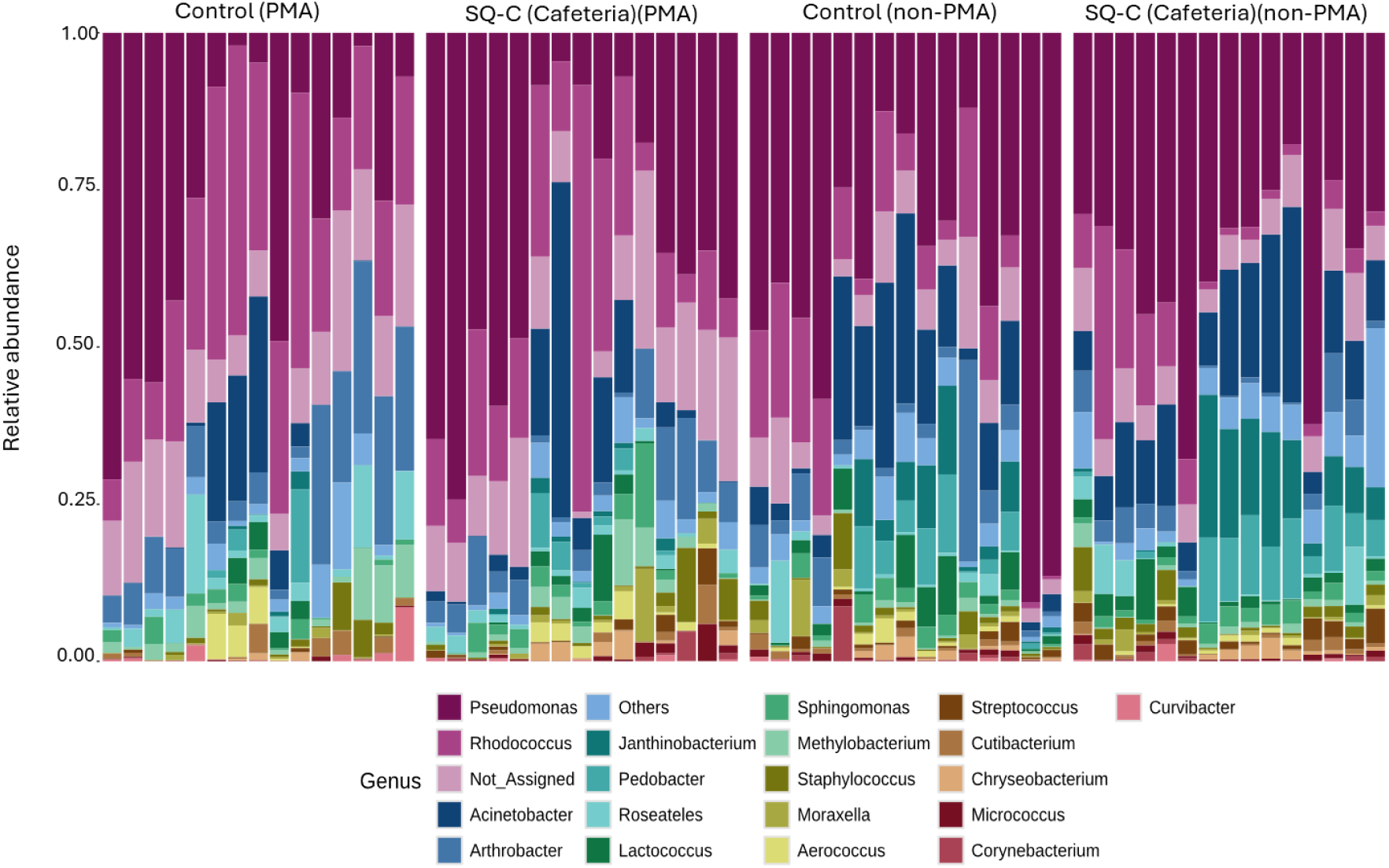
Relative abundance of the top 20 bacterial genera detected on tables in cafeteria. Each stacked bar represents an individual surface sample on either non-PMA or PMA treated control or SiQAC-based (SQ-C) surface with bar height indicating the proportion of each genus within that sample.

**Figure S14.**
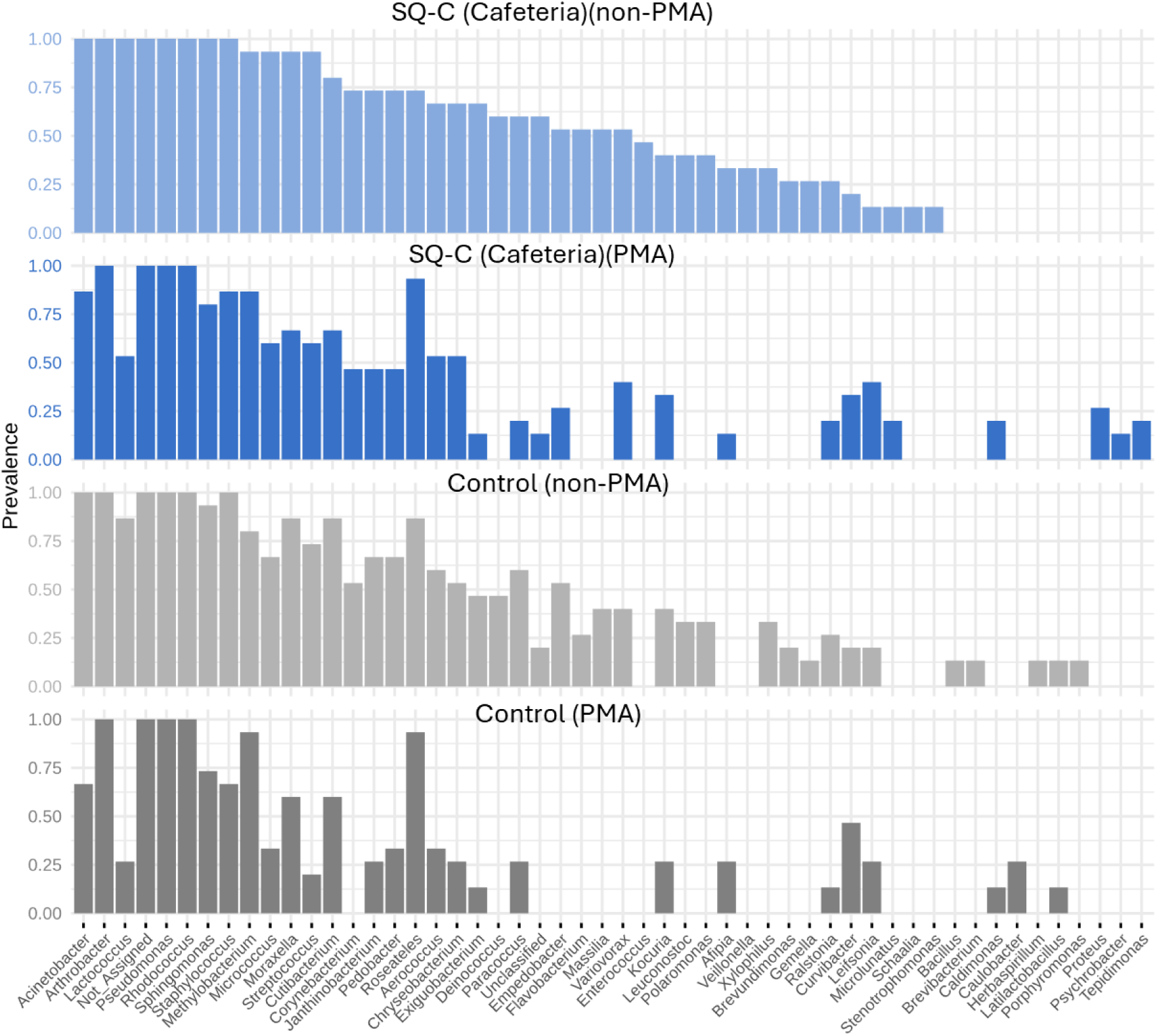
Prevalence of bacterial genera across samples collected from tables in cafeteria. Bars represent the proportion non-PMA or PMA treated control or SiQAC-based (SQ-C) surface samples in which each genus was detected, applying a relative abundance threshold of 0.001 and a sample prevalence cutoff of 10%. Genera are displayed in descending order of prevalence within each condition.

**Figure S15.**
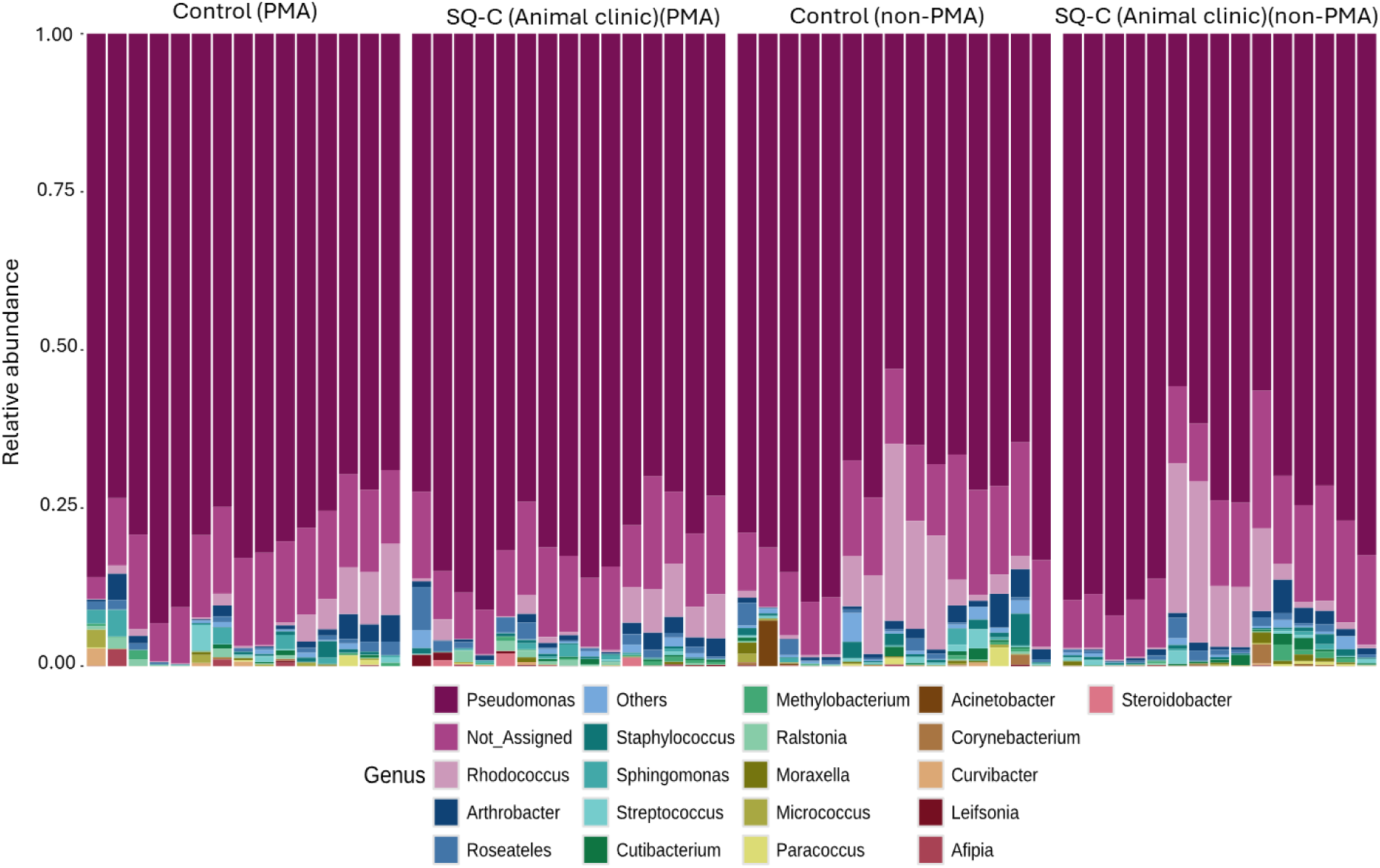
Relative abundance of the top 20 bacterial genera detected on tables in animal clinic. Each stacked bar represents an individual surface sample on either non-PMA or PMA treated control or SiQAC-based (SQ-C) surface with bar height indicating the proportion of each genus within that sample.

**Figure S16.**
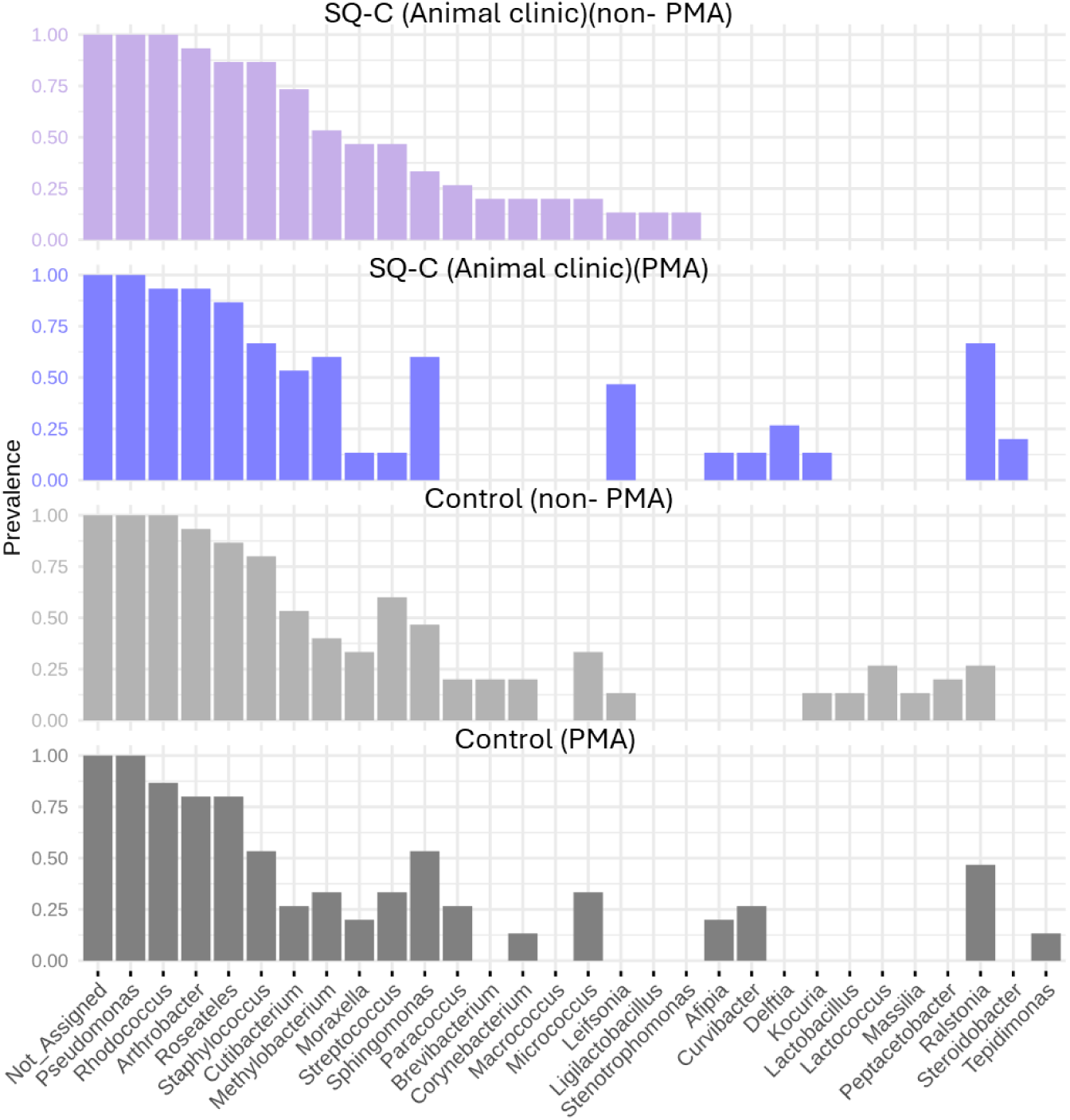
Prevalence of bacterial genera across samples collected from the tables in animal clinic Bars represent the proportion non-PMA or PMA treated control or SiQAC-based (SQ-C) surface samples in which each genus was detected, applying a relative abundance threshold of 0.001 and a sample prevalence cutoff of 10%. Genera are displayed in descending order of prevalence within each condition.

**Table S1.** Relative abundance of genera in empty swab controls. The detected genera can be associated with environmental (*Rhodococcus*, *Gordonia*, *Methylobacterium-Methylorubrum*, *Afipia*, and *Pelomonas)*, reagent (*Renibacterium)*, or human skin (*Cutibacterium*) background contamination. The relatively high abundance of *Renibacterium* empty swab control suggests that this taxon was introduced during the workflow and was not a true biological signal from surface samples.

|  | Relative abundance |
| --- | --- |
| *Renibacterium* | 0.40 |
| *Rhodococcus* | 0.27 |
| *Cutibacterium* | 0.07 |
| *Not_Assigned* | 0.07 |
| *Gordonia* | 0.05 |
| *Methylobacterium* | 0.05 |
| *Afipia* | 0.02 |
| *Pelomonas* | 0.01 |

**Table S2.**
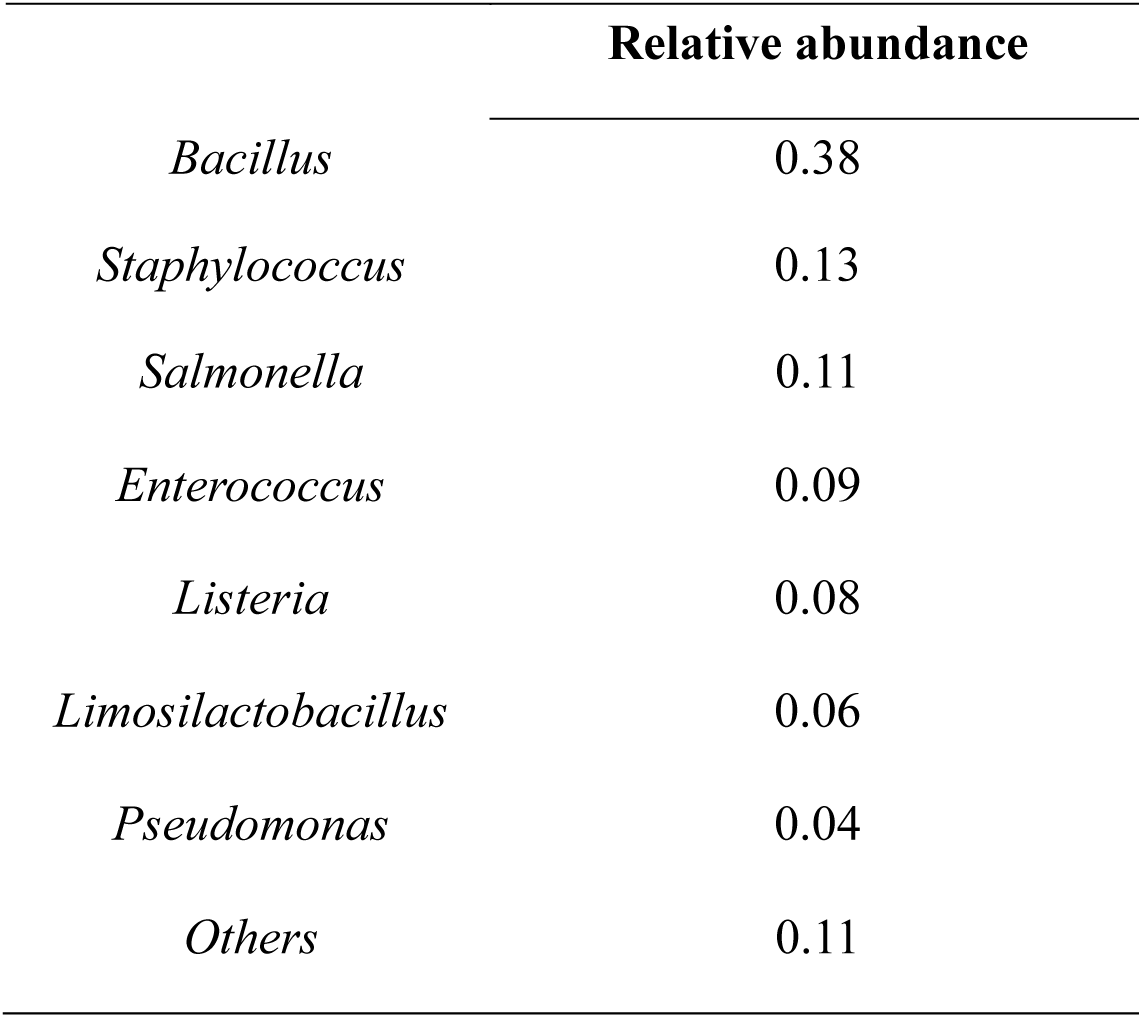
Relative abundance of the genera in the mock community. The used Zymo mock community theoretically consists of 12% of each of those species: *Pseudomonas aeruginosa, Escherichia coli, Salmonella enterica, Lactobacillus fermentum, Enterococcus faecalis, Staphylococcus aureus, Listeria monocytogenes, Bacillus subtilis* and this aligns with the experimental results except *_E_scherichia*, which was likely represented within *Others*.

|  | Relative abundance |
| --- | --- |
| *Bacillus* | 0.38 |
| *Staphylococcus* | 0.13 |
| *Salmonella* | 0.11 |
| *Enterococcus* | 0.09 |
| *Listeria* | 0.08 |
| *Limosilactobacillus* | 0.06 |
| *Pseudomonas* | 0.04 |
| *Others* | 0.11 |

